# Regionally divergent drivers of historical diversification in the late Quaternary in a widely distributed generalist species, the common pheasant *Phasianus colchicus*

**DOI:** 10.1101/2019.12.21.881813

**Authors:** Simin Liu, Yang Liu, Edouard Jelen, Mansour Alibadian, Cheng-Te Yao, Xintong Li, Nasrin Kayvanfar, Yutao Wang, Farhad Vahidi, Jianlin Han, Gombobaatar Sundev, Zhengwang Zhang, Manuel Schweizer

**Author notes:** **Correspondence**: Yang Liu, State Key Laboratory of Biocontrol, School of Life Sciences and Department of Ecology, Sun Yat-sen University, Guangzhou 510275, China.

## Abstract

**Aim:** Historical factors such as Pleistocene climate cycles and associated environmental changes have influenced the phylogeographic structure and demographic dynamics of many species. Resulting patterns not only depend on species’ life-history but also vary regionally. Consequently, different populations of species with large ranges over different biomes might have experienced divergent drivers of diversification and show different population histories. Such a representative species is the common pheasant *Phasianus colchicus*, an ecological generalist with a wide distribution in the Palearctic and at the edge of the Oriental region. We aimed at identifying distinct phylogeographic lineages of the common pheasant and investigating their evolutionary trajectories.

**Study location:** Asia

**Methods:** We used coalescent approaches to describe the phylogeographic structure and to reconstruct the spatio-temporal diversification and demographic history of the common pheasant based on a comprehensive geographic sampling of 265 individuals genotyped at seven nuclear and two mitochondrial loci.

**Results:** The common pheasant diversified during the late Pleistocene into eight distinct evolutionary lineages which only partly correspond to traditional morphological groups. It originated at the edge of the Qinghai-Tibetan plateau and spread from there to East and Central Asia. Only the widely distributed genetically uniform lowland lineage of East Asia showed a recent range and population expansion, starting during last glacial. More phylogeographic structure was found elsewhere with lineages showing no signs of recent range expansions. One lineage of subtropical south-central China this is the result of long-term isolation in a climatically stable and topographically complex region. In others from arid Central Asia and China, demographic and range expansions were impeded by repeated population fragmentation during dry glacial and recent aridification. Given such a phylogeographic structure and demographic scenarios among lineages, we proposed split the range-wide common pheasant into three species.

**Main conclusions:** Spatio-temporal phylogeographic frameworks of widespread species complexes such as the common pheasant provide valuable opportunities to identify regionally divergent drivers of diversification.

## 1 INTRODUCTION

Distributional ranges of free-living organisms, in term of size, shape and boundaries, vary substantially in geographical space (Brown *et al.*, 1996; Newton, 2003). The range of organisms increases on average with increasing latitude, a correlation known as Rapoport’s rule which is chiefly driven by latitudinal range extent (Stevens, 1989). Particularly large ranges are often discontinuous and involve different subspecies, such that breeding range size has been shown to be a good predictor of subspecies richness (Phillimore *et al.*, 2007). In line with this, intraspecific divergence, a proxy for ‘subspeciation’, has been shown by meta-analyses to be generally higher in mammals and nonmigratory birds at higher latitudes in the Northern Hemisphere, with subspecies richness being positively correlated with environmental harshness, which generally increases with latitude (Botero *et al.*, 2014; Weir, 2014). Furthermore, subspecies richness can be higher in species whose ranges include several biomes (Phillimore *et al.*, 2007).

Apart from these different environmental correlates, historic factors also play a role, and subspecies richness can be positively correlated with historical exposure to glaciation (Botero *et al.*, 2014). Recent range dynamics and phylogeographic patterns have been found to be strongly influenced by Pleistocene climate cycles and associated landscape alterations (e.g. Hewitt, 2000; Hewitt, 2004). However, species’ reactions to these environmental changes were manifold and largely depended on their life history, with faster diversification being positively correlated with ecological specialization, resident life-style and low dispersal ability (e.g. Hung *et al.*, 2017). While mesic temperate and boreal species or species complexes showed range contractions during cold dry glaciations and range expansion during warmer wetter interglacial periods (e.g. Hewitt, 2000), arid-adapted species could expand their ranges in ice-free areas during dryer glacial periods (Garcia *et al.*, 2011a; Garcia *et al.*, 2011b; Kearns *et al.*, 2014; Alaei Kakhki *et al.*, 2018).

In contrast to temperate mesic species, the subtropical avifauna of East Asia showed range contraction during interglacial rather than glacial periods, probably caused by sensitivity to climate variation during the latter period (Dong *et al.*, 2017). In addition, interactions between climate change and geographic features such as topography led to regionally different phylogeographic patterns in the Holarctic and adjacent regions. For instance, bird species from mountainous south-central China often show strongly structured but long-term stable populations as the result of a combination of complex mountainous landscapes and a climate which was mildly affected by Pleistocene climate change (Qu *et al.*, 2014; Lei *et al.*, 2015; Cai *et al.*, 2018). As a consequence, different subspecies (reflecting recent population splits) of species with large ranges in East Asia, south-central China and Central Asia might have experienced divergent drivers of diversification and show different evolutionary trajectories. Phylogeographic studies on widespread species occurring in a broad range of ecological and climatic conditions over different biomes can thus provide a valuable opportunity to test for the impact of climatically induced environmental change on lineage diversification and demography (Pyron & Burbrink, 2009; Statham *et al.*, 2014).

Such a representative species is the common pheasant *Phasianus colchicus*, which has a wide distribution throughout the southern part of the eastern Palearctic, occurring across different biomes with an exceptional number of described subspecies (Figure 1). Apart from introduced populations, e.g. in Europe or North America, the common pheasant occurs from the Caucasus over Central Asia and from the eastern edge of the Qinghai-Tibetan plateau to eastern China and south-east Siberia. Its range enters the Oriental region in southern China and northern Vietnam and Myanmar (McGowan & Kirwan, 2019). It occurs from sea level up to 3500 m on the Qinghai-Tibetan plateau in a wide range of climates and habitats. It is an ecological generalist often associated with riverine or lakeside vegetation, woodland edges and scrubby open-habitats, but it avoids dense forests and very dry areas (Madge *et al.*, 2002). Currently, 30 subspecies are usually recognized (McGowan & Kirwan, 2019), which have been divided into five groups primarily based on differences in males’ plumage and to a lesser extent on biogeography: *colchicus* group or ‘black-necked pheasant’ (4 subspecies), *mongolicus* group or ‘Kyrghyz pheasant’ (2 subspecies), *principalis-chrysomelas* group or ‘white-winged pheasant’ (5 subspecies), *tarimensis* group or ‘Tarim basin pheasant’ (2 subspecies) and *torquatus* group or ‘grey-rumped pheasant’ (17 subspecies) (Madge *et al.*, 2002) (McGowan & Kirwan, 2019) (cf. Figure 1).

**Figure 1.**
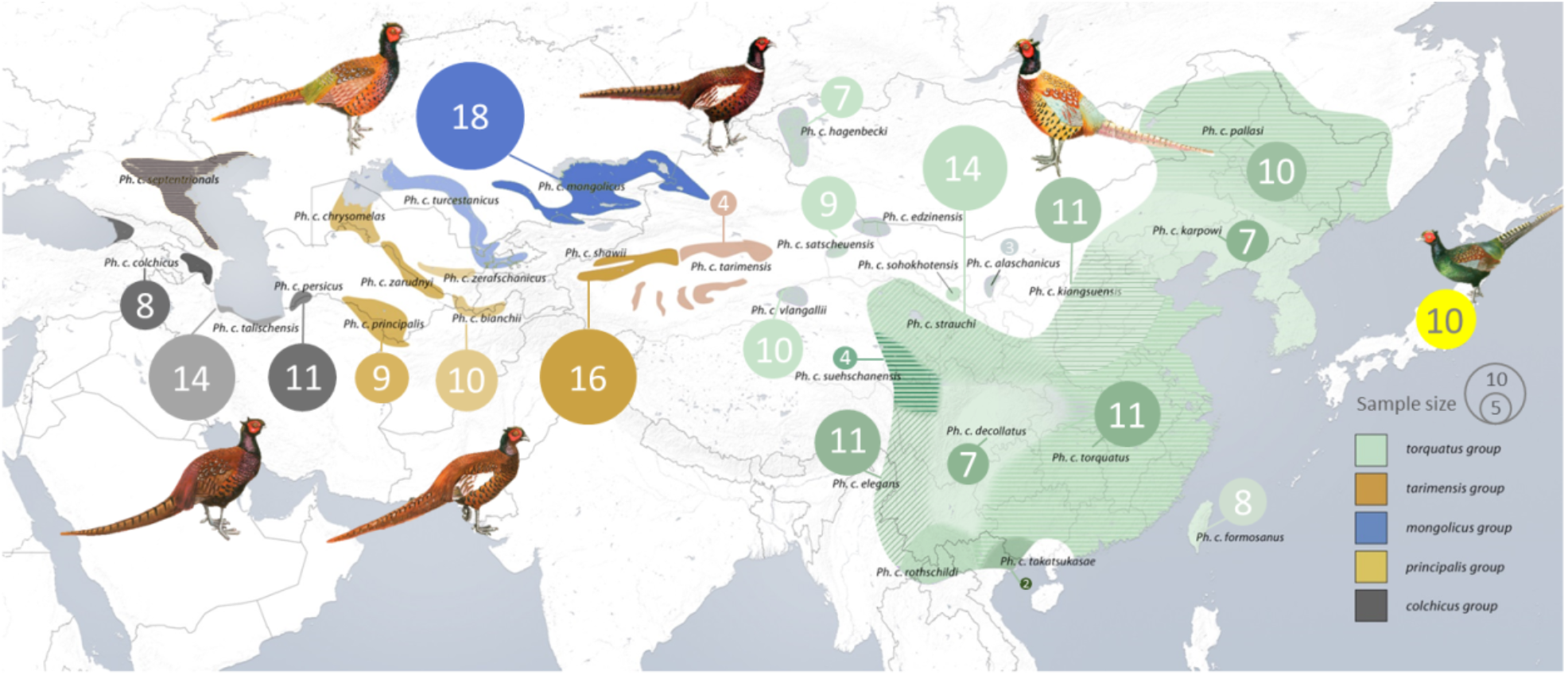
Geographic distribution of the different subspecies of the common pheasant *Phasianus colchicus*. The ranges of the five traditionally defined subspecies groups are shown in different colors: black-necked pheasant or *colchicus* group (dark grey), white-winged pheasant or *principalis-chrysomelas* group (ochre), Kirghiz pheasant or *mongolicus* group (blue), olive-rumped pheasant or *tarimensis* group (purple) and grey-rumped pheasant or *torquatus* group (green). The green pheasant *Phasianus versicolor* from Japan was selected as outgroup (yellow). The size of each circle indicates the number of samples used in this study.

Males of both the *principalis-chrysomelas* and the *mongolicus* groups are characterized by white wing-coverts, with the former being more orange-yellowish instead of coppery-colored on the upperparts. The other three groups display buffy wing-coverts with the rump being reddish-brown in the *colchicus* group, green with a yellowish cast in the *tarimensis* group and green with greyish or blue cast in the *torquatus* group. Complete white collars are found in the *torquatus* and *mongolicus* group, while they are only partially developed in the *principalis-chrysomelas* group, with the exception of the subspecies *zeravshanicus*, which has a well-developed collar. Within the groups, morphological differences among subspecies are minor and mainly concern coloration of different plumage parts and appear often clinal when subspecies’ ranges are contiguous (Madge *et al.*, 2002).

A recently published study using two mitochondrial DNA (mtDNA) fragments and two autosomal nuclear introns revealed incongruence between these morphology-based subspecies groups and phylogenetic relationships (Kayvanfar *et al.*, 2017). The subspecies *P. c. elegans* from the Hengduan mountains (south-central China) and northern Myanmar, which is traditionally included in the *torquatus* group, was found to be the sister group of all remaining taxa, although with low support. Moreover, the remaining taxa in the *torquatus* group formed no monophyletic clade either. A well-supported clade containing all other groups was nested within the taxa of the *torquatus* group. Within this clade, the monophyly of the *colchicus*, *mongolicus* and *principalis-chrysomelas* groups received robust support, but the study suffered from low sample size with limited geographic coverage.

We therefore aimed here at building a comprehensive understanding of the evolutionary history of the common pheasant. To test the phylogeographic patterns indicated by Kayvanfar *et al.* (2017) we increased the taxon sampling by a factor of more than three with a larger coverage from the species’ entire range in Asia, and genotyped these individuals at six nuclear introns, one Z-linked and two mtDNA loci. We used coalescent approaches to reconstruct the spatio-temporal diversification and demographic history of the common pheasant’s different evolutionary lineages to test for regionally different impacts of past climate and associated environmental change.

## 2 MATERIALS AND METHODS

### 2.1 Sample preparation

Blood, muscle or feather samples of 204 individuals of common pheasants were collected from 90 locations throughout the species’ geographical range, representing 22 out of the recognized 30 subspecies (Figure 1, Table S1). Birds from captivity were sampled for one taxon (*bianchii*). We used the green pheasant *Phasianus versicolor* (n (number of samples) = 10) endemic to Japan as outgroup. We analysed two mitochondrial markers, six autosomal nuclear introns and one Z-linked nuclear intron (Figure 2, Table S2-S3). See Supporting information for laboratory protocol and sequence editing.

**Figure 2.**
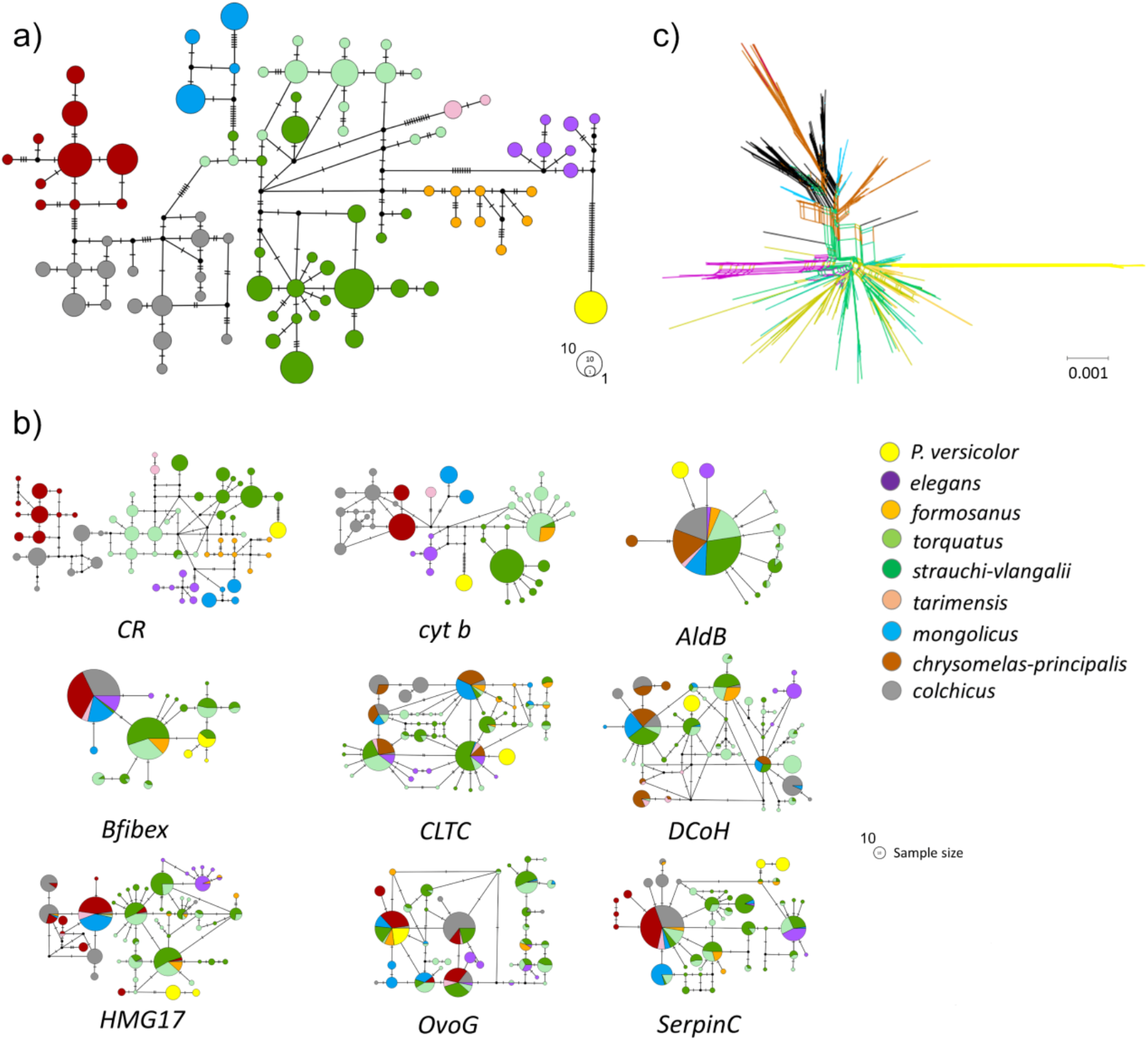
Common pheasant haplotype networks of a) the two mtDNA loci cytochrome b and control region, and b) of the seven nuclear introns. Size of circles is proportional to the number of sampled haplotypes or alleles, colors correspond to the different evolutionary lineages. Dashed lines indicate inferred unsampled haplotypes. c) Neighbor-net network of the nuclear dataset. Colors of branches correspond to the different evolutionary lineages.

### 2.2 Phylogeographic, demographic and migration analyses analysis

A detailed description of methods is available in Supporting Information. To visualize genetic variation, a median-joining haplotype networks was constructed for each marker in PopART v4.8.4 (Leigh & Bryant, 2015). Furthermore, we generated a neighbor-net phylogenetic network based on all nuclear loci in SplitsTree v4.14.4 (Huson & Bryant, 2006). The approach of Li *et al.* (2010) was followed to estimate substitution rates of all loci. The ration of mean genetic distances of each marker and the mean of cytochrome b (cyt b) was multiplied by average substitution rate of the latter for Galliformes (0.0238 substitutions/site/million year: (Weir & Schluter, 2008). A dated gene tree based on the mtDNA dataset was reconstructed in BEAST v2.4.7 (Bouckaert *et al.*, 2014). Based on concatenated dataset of two mtDNA markers and the alleles of six nuclear introns, a calibrated species tree was reconstructed using *BEAST v2.4.7 (Bouckaert *et al.*, 2014). The locus AldB located on the Z chromosome was excluded from this analysis as information on sex was not available for all individuals. Based on the results of the mtDNA gene tree estimated with BEAST (see below), samples of eight different evolutionary lineages within common pheasant were set as ‘taxa’. *BEAST was moreover used to reconstruct the colonization routes of the eight evolutionary lineages based on the concatenated dataset as above using GEO_SPhERE and BEAST-CLASSIC packages (Lemey *et al.*, 2009).

Changes in effective population size (*Ne*) through time in the different evolutionary lineages (see below) were reconstructed with the Extended Bayesian Skyline Plot (EBSP) by setting up the coalescent extend Bayesian Skyline as tree prior in BEAST based on the concatenated dataset of all markers. Due to a low sample size, no inference of demographic history was performed for the *tarimensis* and the *formosanus* groups.

Since EBSP analyses assume no gene flow after divergence between lineages, we additionally inferred non-equilibrium scenarios between parapatric lineages based on an isolation-with-migration model. This allowed us to assess migration rates, effective population size and divergence time. Analyses were run on each pair of parapatric lineages using IMa2 (Hey, 2010a, b).

### 2.3 Species delimitation

We moreover conducted a species delimitation analysis in BPP v.3.4 (Yang, 2015) using the species-tree of the *BEAST analyses (see below) as the user-specified guide tree (Yang & Rannala, 2010; Rannala & Yang, 2013) treating the eight distinct evolutionary lineages within common pheasant as potential species with equal prior probabilities to all potential species delimitation models. More information on parameter settings is given in Supplementary Information.

## 3 RESULTS

### 3.1 Sequence characteristics

We obtained an alignment of 204 individuals, including 1188 bp of two mitochondrial loci and 4421 bp of seven introns. No locus deviated significantly from neutrality. Detailed information including sequence length, standard diversity indices, best-fitting models of nucleotide substitution and substitution rates are shown in the supporting information (Table S4-S5).

### 3.2 Phylogeographic analyses

In all BEAST analyses, parameters converged among different runs. After combining the three independent runs with 10% burnin each, ESS values were >200 for all except the proportion of invariant sites in the substitution model for cytochrome b (ESS ≥180). In the resulting maximum clade credibility mtDNA eight major lineages were revealed within common pheasant and their monophyly robustly supported, except in lineages 4 and 7 (Figure 3). While the green pheasant diverged in the Mid-Pleistocene, diversification within common pheasants started at the beginning of the Late Pleistocene around 0.7 million years ago (Ma) and divergence of the eight major evolutionary lineages continued until the last glacial period (see Figure 3 for mean node ages and 95% highest posterior density HPD intervals).

**Figure 3.**
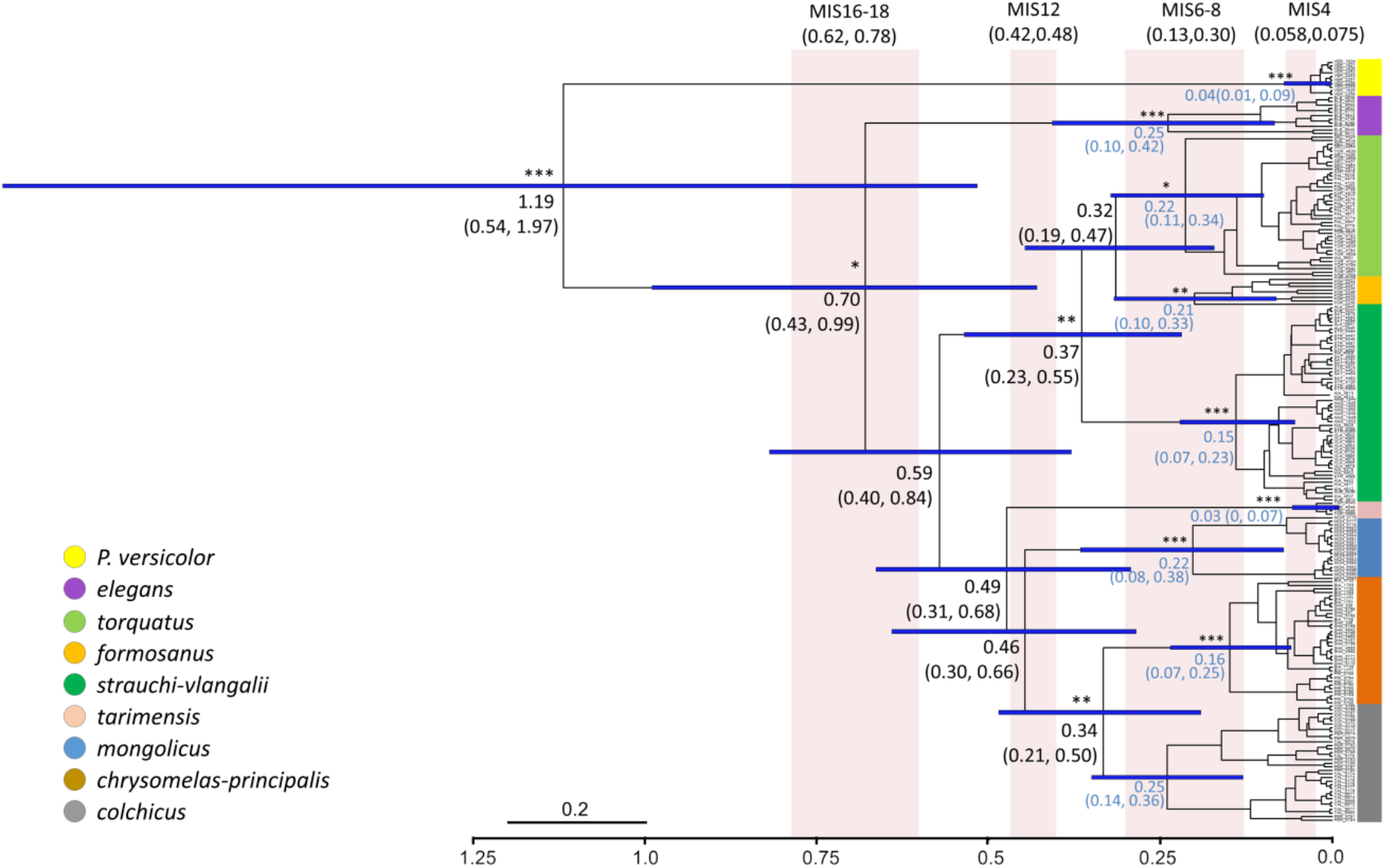
Time-calibrated maximum clade credibility mtDNA gene tree (two mitochondrial regions, control region and cytochrome b gene, 1188bp totally) reconstructed in BEAST. Mean ages and their 95% highest posterior density intervals are given at nodes. The blue bars illustrate the 95% highest posterior density intervals of inferred node ages. Asterisks at nodes represent different levels of posterior node probabilities (*** = 1.0, ** > 0.9 and * >0.8). Pink bars indicate the main glacial periods of the late Pleistocene.

Lineage 1 (*elegans* group) comprised all samples of the taxon *elegans*. It formed the sister group to all remaining common pheasant taxa, although this position was not robustly supported. The remaining taxa were split in two clades, one containing samples of all taxa from Central Asia, the other all from East Asia, but only the latter received robust support. The samples of all East Asian common pheasant were further split into three clades, whose sequence of lineage splitting was not robustly supported. Lineage 2 (*strauchi-vlangalii* group) contained samples of *alaschanicus*, *hagenbecki*, *kiangsuensis*, *satscheuensis*, *suehschanensis*, *strauchi* and *vlangalii*. Lineage 3 (*formosanus* group) comprised all samples of *formosanus* from Taiwan. The monophyly of Lineage 4 (*torquatus* group) was not robustly supported, contained samples of *decollates*, *karpowi*, *pallasi*, *takatsukasae* and *torquatus*. One individual each from the area of the taxa *kiangsuensis*, *strauchi* and *suehschanensis* of the *strauchi-vlangalii* group clustered within the *torquatus* group. In the Central Asian clade, lineage 5 (*tarimensis* group), containing all samples of *tarimensis*, was revealed as sister group of the remaining taxa, although not robustly supported. Also the position of lineage 6 (*mongolicus* group), containing all samples of *mongolicus*, was not robustly supported. The two remaining main clades of Central Asian taxa clustered robustly together. Lineage 7 (*principalis-chrysomelas* group) included samples of *bianchii*, *principalis* and *shawii*. Lineage 8 (*colchicus* group) comprised the taxa *colchicus*, *persicus* and *talischensis*, but its monophyly was not robustly supported. The eight evolutionary lineages were also reflected in the median-joining networks of the two mtDNA markers. The nuclear introns instead showed no separation between the eight lineages and a lot of allele sharing (Figure 2).

In the *BEAST analyses, there was good convergence among parameters of the four independent runs. After combining the four runs with 10% burnin each, ESS values were >200 for all parameters except the tree likelihood of *SerpinC* (ESS=180). The topology of the *BEAST maximum clade credibility species tree among the eight predefined major evolutionary lineages had no conflict in supported nodes with the BEAST mtDNA gene tree, except that the *tarimensis* group and the *principalis-chrysomelas* group formed a strongly supported clade, instead of *principalis-chrysomelas* with the *colchicus* group. (Figure 4). The following relationships were not robustly supported: sister group position of the *elegans* group to the remaining common pheasants, monophyly of the East Asian lineages, and the sister group position of the *mongolicus* group to the remaining Central Asian lineages. Unlike in the BEAST mtDNA gene tree, a clade comprising the *strauchi-vlangalii* and *torquatus* group was robustly supported. The divergence of the green pheasant was dated at around 0.6 Ma (see Figure 4 for mean node ages and 95% HPD intervals). Diversification within common pheasants was revealed at around 0.2 Ma, while the split between Central Asian and East Asian lineages was estimated at around 0.13 Ma.

**Figure 4.**
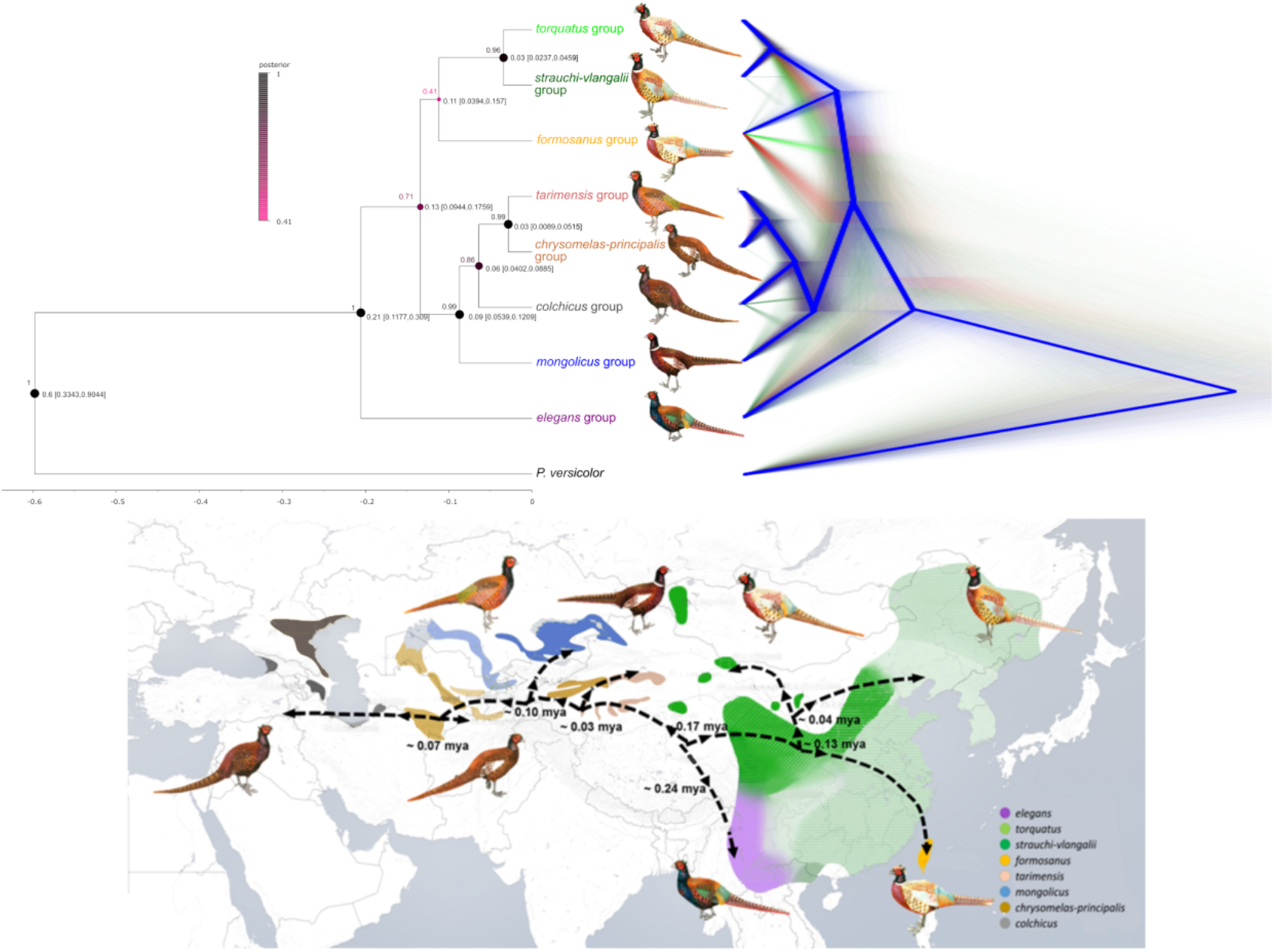
Spatio-temporal lineage divergence of common pheasant. Top left: Time calibrated maximum clade credibility species tree reconstructed in *BEAST based on two mtDNA markers and six nuclear introns. Numbers at nodes represent mean node ages with 95% highest posterior density intervals. Posterior probabilities of nodes are illustrated by a color code. The blue bars represent 95% highest posterior density distributions for node ages. Top right: Cladogram showing the uncertainty of the sampled topologies over the posterior distribution. Different topologies from the posterior distribution are shown in different colors. Blue: Most popular topologies. Red: 2nd most popular topologies. Pale green: 3rd most popular topologies. Dark green: 4th most popular topologies. Bottom: Reconstruction of colonization routes of the common pheasant with *BEAST and SPREAD. Arrows indicate the direction of colonization; mean divergence times are additionally shown.

The reconstructed colonization routes of common pheasants revealed three colonization directions from the edge of the Qinghai-Tibetan plateau: 1) to the south-east towards around 0.23 (0 – 0.35) Ma resulting in the *elegans* group, and almost simultaneously to 2) East Asia and 3) Central Asia around 0.17 (0.12 – 0.23) Ma (Figure 4).

### 3.3 Demographic and migration analyses

According to EBSP analyses, a stable population size is only rejected for the *torquatus* group, with a median for the number of population size change events and a with a 95% HPD interval ranging from one to three. The 95% HPD interval included zero for all other lineages and stable population sizes could not be rejected. Accordingly, a population size change in the *torquatus* group was indicated in the EBSP plot with an increase in effective population size during the last 50,000 years (Figure 5). The effective population sizes of the other groups seemed basically stable in the EBSP plots, with only slight apparent recent decrease in the *mongolicus* group and slight recent increase in the *strauchi-vlangalii* group.

**Figure 5.**
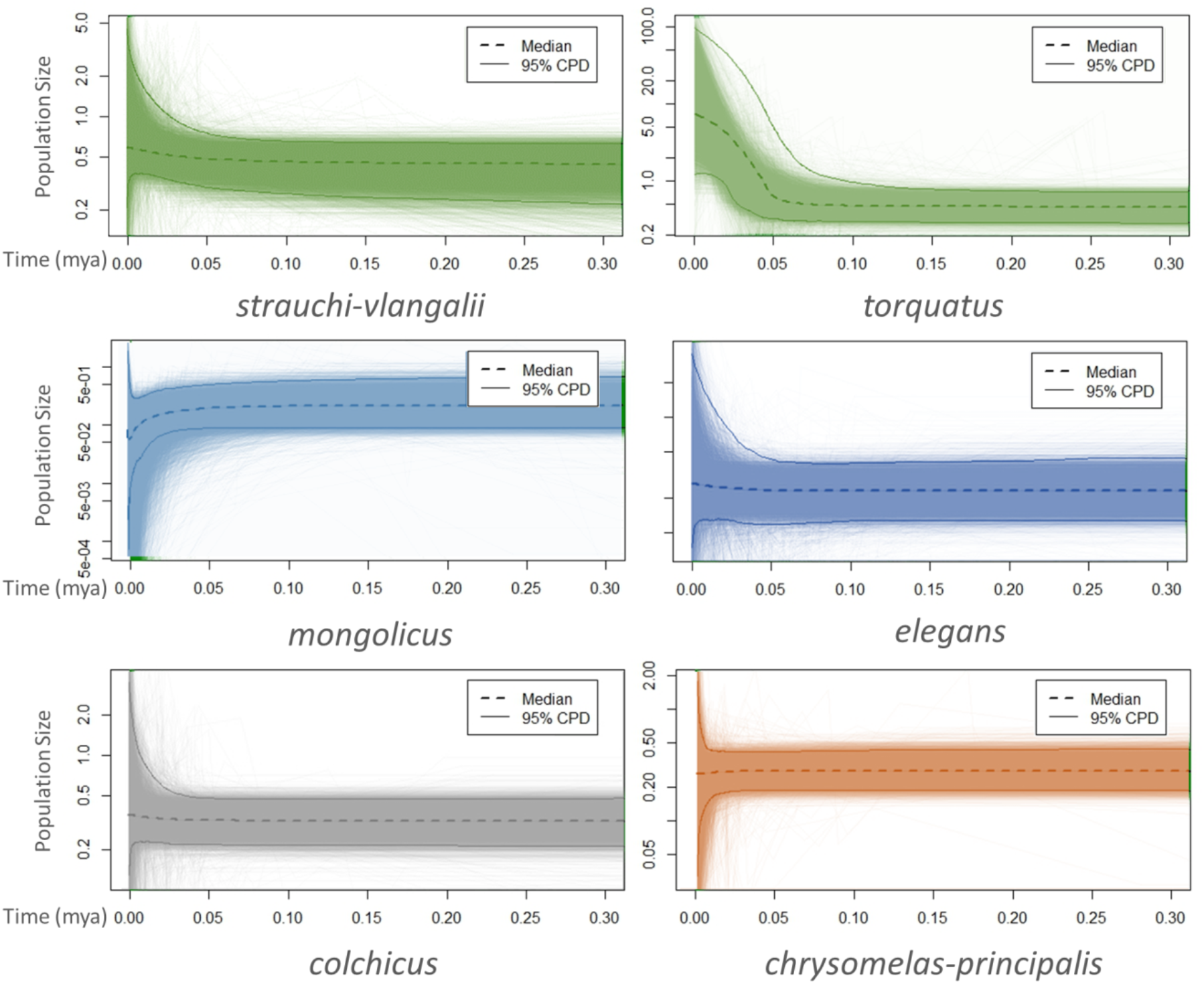
Extended Bayesian Skyline Plots based on two mtDNA markers and seven nuclear introns estimated in BEAST for six different evolutionary lineages of the common pheasant. Time scale is in millions of years before present. The dashed lines show median values, shaded colors correspond to the 95% highest posterior density interval.

Gene flow significantly differing from zero among parapatric groups was only found in the *elegans* to the *strauchi-vlangalii* group, from the *torquatus* to the *strauchi-vlangalii* group, from the *formosanus* to the *strauchi-vlanglaii* group, from the *strauchi-vlangalii* to the *tarimensis* group, between the *strauchi-vlangalii* and *mongolicus* groups, between the *tarimensis* and *principalis-chrysomelas* groups and from the *principalis-chrysomelas* to the *mongolicus* group (Figure 6). Values for the highest probability of the posterior distribution for the population migration rates 2NM were <1 except from the *torquatus* to the *strauchi-vlangalii* group and from the *formosanus* to the *strauchi-vlanglaii* group. Effective populations sizes were generally highest in the *torquatus* and *strauchi-vlangalii* groups and lower in the others, with lowest values for the *mongolicus* and *tarimensis* groups (Figure S1). Posterior distributions of divergence time estimates between parapatric populations were not unimodal in several cases (Figure S2). However, estimates with highest posterior density were generally between those of the BEAST and the *BEST analyses.

**Figure 6.**
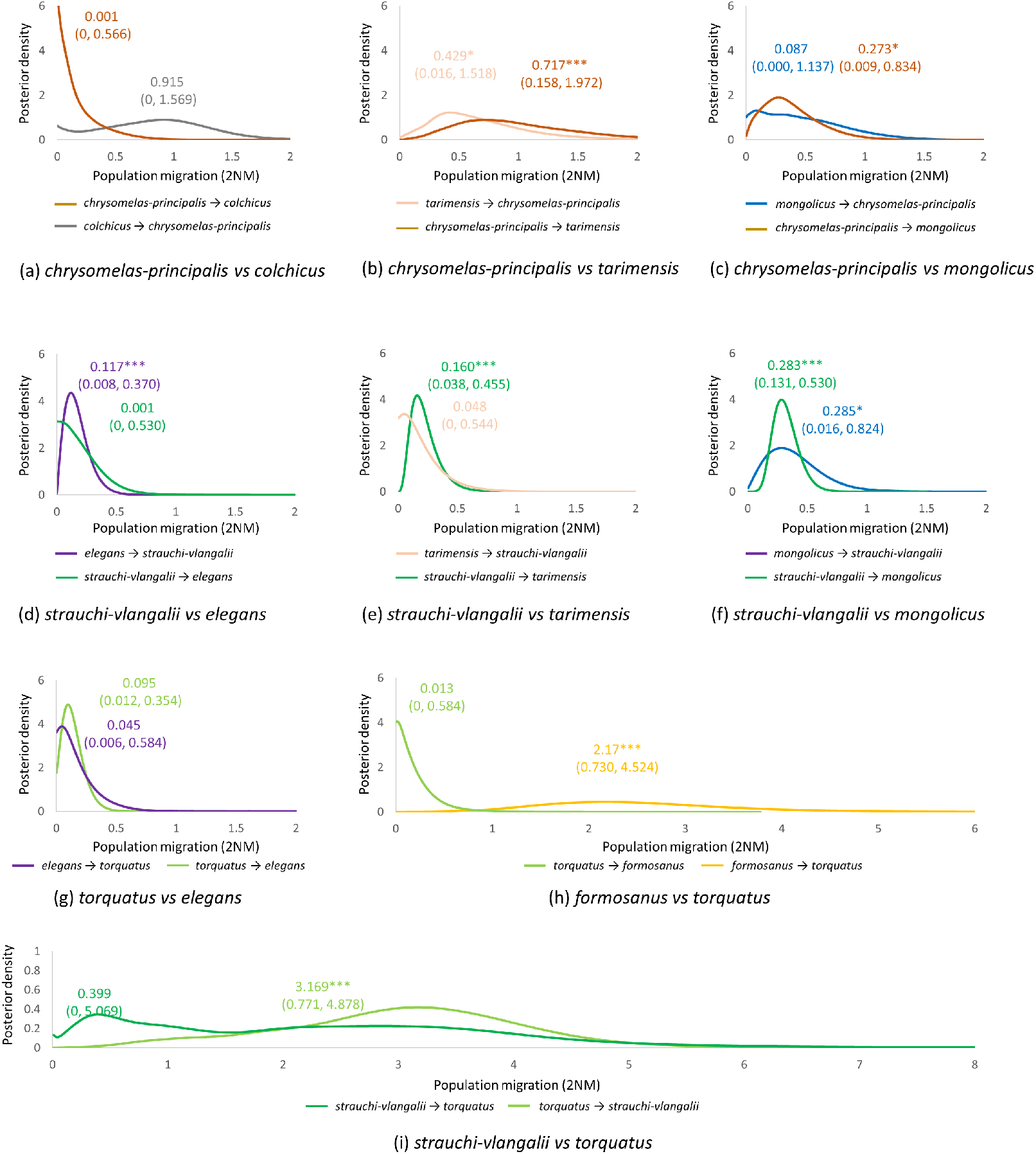
Posterior probability distribution of population migration between parapatric populations of the evolutionary lineages of the common pheasant as estimated with IMa2 based on two mtDNA markers and seven nuclear introns. Values indicate the highest values of the posterior distribution and the 95% highest posterior density interval. Asterisks represent different levels of significant differences of migration parameters from zero (p < 0.001 (***), p < 0.01 (**) and p < 0.05 (*)).

### 3.4 Species delimitation

In all species delimitation analyses conducted in BPP using different prior settings for the population size and divergence time parameters, the eight different evolutionary lineages of the common pheasant were delimited as eight different species with posterior probabilities of one.

## 4 DISCUSSION

### 4.1 Phylogeographic structure: eight distinct evolutionary lineages

We revealed the common pheasant to consist of eight distinct evolutionary lineages which only partly corresponded to traditionally recognized subspecies groups defined chiefly on morphological similarities. Five of these eight lineage lineages occur in the arid parts at the edge of the Qinghai-Tibetan plateau as well as in Central Asia, while no phylogeographic structure was revealed over a large area from south-east China to the Amur and Ussuri region in south-east Siberia.

In congruence with Kayvanfar et al. (2017), the subspecies *P. c. elegans* of the Hengduan mountains of south-central China and northern Myanmar was found to be the sister group of the remaining common pheasant taxa. It has traditionally been included in the *torquatus* group. The taxon *rothschildi* from south-east Yunnan in southern China and north-western Vietnam was not sampled here but is morphologically similar to *elegans* (Madge *et al.*, 2002), so further genetic studies are needed to show if it is part of this basal clade. Divergence within the remaining populations of the traditional *torquatus* group was indicated by previous studies with limited sampling based on mtDNA (Qu *et al.*, 2009; Liu *et al.*, 2010; Qu *et al.*, 2017), and we found them to be split in three distinct evolutionary lineages. 1) The *strauchi-vlangalii* group (*containing alaschanicus*, *hagenbecki*, *kiangsuensis*, *satscheuensis*, *suehschanensis*, *strauchi* and *vlangalii*) is characteristic of comparatively arid areas of the Loess plateau in northern China, but also occurs on the north-eastern part of the Qinghai-Tibetian plateau south to northern Sichuan, with scattered populations in the Qaidam Basin, Nei Mongolia and Western Mongolia (McGowan & Kirwan, 2019). The taxa *edzinensis* and *sohokhotensis* of central-northern China, not sampled here, most probably belong to this lineage (cf. Figure 1). 2) The *torquatus* group (containing *decollates*, *karpowi*, *pallasi*, *takatsukasae* and *torquatus*) occurs from the humid subtropical parts of south and eastern China to northern China, extreme south-east Siberia and the Korean peninsula. 3) The *formosanus* group comprising all samples from Taiwan formed an additional distinct evolutionary lineage.

The four distinct evolutionary lineages revealed in Central Asia corresponded to the traditionally defined groups, i.e. *tarimensis*, *mongolicus*, *principalis-chrysomelas* and *colchicus*. The taxon *shawii* replaces *tarimensis* in the western part of the Tarim basin in western China, and was traditionally included in the *tarimensis* group given that the Tien Shan range separates it from populations of other Central Asian groups. However, it shares morphological features such as whitish wing-coverts and reddish rump and upper-tail coverts with birds of the *principalis-chrysomelas* group, and we unequivocally found it to cluster within the *principalis-chrysomelas* group. Within the *principalis-chrysomelas* group we only had samples of the southern taxa *principalis* and *bianchii*; the more northerly taxa *chrysomelas*, *zerafschanicus* and *zarudnyi*, traditionally included in this group, were unsampled. We only had samples of the nominate subspecies of the *mongolicus* group and could not include the southern taxon *turkestanicus* with an adjacent distribution to taxa of the *principalis-chrysomelas* groups in eastern Uzbekistan and in the Aral Sea region. Within the *colchicus* group, the taxon *septentrionalis*, which occurs north of the Caucasus in the area of the lower Volga, was not included. Overall fine-scale phylogeographic structure still requires further investigations.

### 4.2 Temporal diversification during the Pleistocene

Diversification within common pheasant started after the Early-Middle Pleistocene Transition between 1.2–0.8 Ma, when climatic cycles increased in duration and amplitude (Maslin *et al.*, 2014). While the initial split was dated at around 0.7 Ma in the mtDNA gene tree, considerably younger divergence events were revealed in the species tree analyses, with diversification beginning at around 0.2 Ma. However, mtDNA coalescence events as estimated in the BEAST gene tree analyses are expected to overestimate divergence dates and pre-date lineage splits estimates with multilocus species tree approaches such as *BEAST (Edwards & Beerli, 2000), and similar discrepancies have been found in other studies (e.g. Drovetski *et al.*, 2015; Kamp *et al.*, 2019). Moreover, gene flow among lineages might confound divergence time estimates in *BEAST, making them appearing more recent. Accordingly, isolation with migration analyses among parapatric evolutionary lineages revealed generally intermediate divergence dates between the time-calibrated mtDNA gene tree and the species tree approach. Given this uncertainty and the rather large confidence intervals of estimated node ages, we cannot relate splits among evolutionary lineages to particular events during the Pleistocene, i.e. glacial or interglacial periods. Our divergence time estimates are considerably younger than that of Kayvanfar *et al.* (2017), who dated the split between *P. colchicus* and *P. versicolor* at about 4 Ma. However, this discrepancy might have been caused by methodological issues. The estimate of the same split by Stein *et al.* (2015) based on fossil calibration was 3 Ma, while that of Cai *et al.* (2018) again based on fossil calibration was more in line with our result. Also the divergence among the *strauchi-vlangalii* and the *torquatus* group estimated by Liu et al. (2010) is in line with our temporal framework.

### 4.3 Spatial diversification: out of the Qinghai-Tibetan plateau and regionally contrasting demographic histories

The common pheasant seems to have originated at the eastern edge of the Qinghai-Tibetan plateau or the adjacent south-west or central Chinese mountains. The Sino-Himalayan region is a diversity hotspot and a diversification center of pheasants in general (Cai *et al.*, 2018). While the avifaunal assemblages of the Himalayas are generally the result of immigration (Johansson *et al.*, 2007; Päckert *et al.*, 2012; Price *et al.*, 2014), the south-west and central Chinese Mountains were revealed as a species-pump in other studies (Päckert *et al.*, 2015; Liu *et al.*, 2016).

From this cradle of diversity at the eastern edge of the Qinghai-Tibetan plateau, the common pheasant expanded its range basically in three directions: 1) to the south-east, leading to the elegans group, and later 2) to the east, leading to the *torqutus*, *strauchi-vlangalii* and *formosanus* groups, and 3) to the west, resulting in the evolution of the Central Asian groups (*tarimensis*, *mongolicus*, *principalis-chrysomelas* and *colchicus* groups). A similar scenario has been proposed by others based on morphology and biogeography (Delacour, 1951; Solokha, 1994) or on limited genetic data (Kayvanfar *et al.*, 2017). Colonization from the Tarim Basin to areas further west might have occurred via a northern route north of the Tien Shan leading to formation of the *mongolicus* group, and via a southern dispersal route south of the Tien Shan along the Pamir-Alai, leading to the formation of the *principalis-chrysomelas* and *colchicus* groups. However, this remains highly speculative, not at least because phylogenetic relationships among Central Asian groups were not unambiguously resolved. Central Asian populations are now chiefly restricted to river valleys, surroundings of larger waterbodies and oases, so riverbeds might have acted as dispersal corridors during colonization towards the west (Solokha, 1994). Until the late Pleistocene, dispersal would have even been possible to the Caspian along ancient river systems crossing today’s Karakorum desert. The Amu Darya (Amo River) then reached the Caspian Sea via the Uzboy River (Leroy *et al.*, 2007).

The *elegans* group occurs in the biodiversity hotspots of the south-west Chinese mountain ranges and northern Myanmar (Myers *et al.*, 2000; Huang *et al.*, 2010; Lei *et al.*, 2015). There, the Hengduan Mountains are considered as an ‘evolutionary powerhouse’ (Zhao *et al.*, 2007; Huang *et al.*, 2010) and their climate has remained relatively stable and warm during the glacial periods of the Pleistocene (Owen *et al.*, 2008). Consequently, the area served as refugia with low extinction risk for many taxa during this time period (Qu *et al.*, 2014; Lei *et al.*, 2015; Xing & Ree, 2017; Cai *et al.*, 2018). However, the topographically complex mountains might have also acted as barriers to dispersal, further promoting lineage diversification (Lei *et al.*, 2015) or impeding range expansion. Accordingly, no change in effective population size was found in the *elegans* group, and it might have evolved in isolation maintaining a stable population size throughout the late Pleistocene.

One of the eastern lineages, the *torquatus* group, is the only common pheasant clade which showed a clear sign of an increase in effective population size towards the present time. This increase started during the last glacial period, which seemingly had no negative impact. Unlike temperate and boreal species in Europe and North America, which experienced range contractions into refugia during glacial periods (Hewitt, 2000; Weir & Schluter, 2004), several subtropical East Asian bird species occurring at different altitudes seem to have expanded their populations during the relatively mild last glacial period after a contraction during the last interglacial period when the climate was highly variable (Dai *et al.*, 2011; Dong *et al.*, 2017; Zhao *et al.*, 2019). After the last glaciation a range expansion might have been possible to areas further north to north-east China and to south-east Siberia, leading to a continuous increase in effective populations. This increase was probably associated with an expansion towards the range of the *strauchi-vlangalii* group, resulting in a suture zones along the eastern edge of the Loess Plateau (Liu *et al.*, 2010). Gene flow of the *torquatus* into the *strauchi-vlangalii* group as indicated in the isolation-with-migration model of the IMa2p analyses, and the presence of mtDNA haplotypes characteristic of the former found in the range of the latter is consistent with this scenario. Liu *et al.* (2010) proposed that these two lineages became differentiated along the eastern edge of the Loess Plateau when the climate became colder and dryer at around 0.24-0.22 Ma. Although aridification as a driver of divergence seems likely, population divergence may not necessarily have happened along the current contact zone.

The insular subspecies *formosanus* forms its own lineage, which is restricted to Taiwan. This island was repeatedly connected to Chinese mainland during cold periods in the Pleistocene, including the last glacial (Li *et al.*, 2010, Wang *et al.* 2019), which could have allowed recurrent gene flow after an initial colonization. We only found signs of gene flow from Taiwan back to mainland China (into the *torquatus* group), but gene flow in the opposite direction might have been masked by the effects of genetic drift in the small island population.

No changes in effective population sizes through time were found in the evolutionary lineages of the common pheasant in the arid parts of the edge of the Qinghai-Tibetan plateau as well as in Central Asia (*strauchii-vlangalii*, *mongolicus*, *principalis-chrysomelas*) and around the Caspian Sea (*colchicus* groups). They may have increased their ranges under warmer and moister conditions of the short interglacials in the late Pleistocene, leading to secondary contact. During the long colder and dryer glacials, range contractions might have led to population isolation. Whether this happened repeatedly, or only during the last glaciation cycle as indicated by the *BEAST multispecies coalescent analyses, needs to be further investigated. While climate in Central Asia as well as northern and western China ameliorated in the beginning of the Holocene after the last glacial maximum, thus there was an abrupt shift back to steppe and desert vegetation between 4 and 6 kya BP (Zhao *et al.*, 2017). This may have hindered population expansion and resulted in the scattered distribution patterns among the different subspecies within the different groups. Based on analyses of plumage features, ancient hybridization between the *mongolicus* and *principalis-chrysomelas* group was suggested (Solokha, 1994). Although we lacked samples of the adjacent populations (see above), we detected low levels of gene flow between *principalis-chrysomelas* and *tarimensis* and from *principalis-chrysomelas* to the *mongolicus* group. However, more extensive and probably bidirectional gene flow could be expected if adjacent populations were sampled in the latter two.

## 5 Conclusion

A phylogeographic framework based on multilocus genetic data of the common pheasant enabled us to test for regionally varying drivers of diversification in this widespread ecological generalist. Lineage diversification and population histories in the eight distinct evolutionary lineages were shaped by regionally varying effects of past climate and associated environmental change in combination with topographic features constraining dispersal. The continuously distributed lowland populations of East Asia (*torquatus* group) are genetically comparatively uniform and might be the result of recent range expansions starting during last glacial period. The remaining range is not only geographically but also genetically more structured as the result of long-term isolation in a climatically stable, topographically complex region (*elegans* group), or caused by the effects of repeated population fragmentation during arid glacial periods or recent aridification (*strauchi-vlanagalii*, *tarimensis*, *chrysomelas-principalis*, *mongolicus* and *colchicus* groups). The remaining evolutionary lineage is a currently isolated island endemic (*formosanus* group).

Given that we detected gene flow among different evolutionary lineages despite adjacent subspecies not being sampled, we refrain from proposing to treat all eight evolutionary lineages as species-level taxa as implied by species delimitation analyses. Integrating phylogeographic structure with morphological variation and taking into account the lineages’ comparatively young ages (cf. Price, 2008), the evolutionary diversification is better reflected by splitting the common pheasant into three species-level taxa: Yunnan Pheasant *P. elegans*, Chinese Pheasant *P. vlangalii* (including the *torquatus*, *strauchi-vlangalii* and *formosanus* groups) and Turkestan Pheasant *P. colchicus* (including *tarimensis*, *chrysomelas-principalis*, *mongolicus* and *colchicus* groups).

Overall, our results demonstrate that spatio-temporal phylogeographic frameworks of ecologically rather uniform widespread species complexes such as the common pheasant indeed provide a valuable opportunity to identify regionally varying drivers of diversification.

## Supporting information

See Supporting information for laboratory protocol and sequence editing.

## ACKNOWLEDGEMENTS

Computational machinery time work was granted by Special Program for Applied Re-search on Super Computation of the NSFC-Guangdong Joint Fund (the second phase) under Grant No. U1501501 to Yang Liu. This study was supported by National Science Foundation of China to YL (No. 31572251). We thank the following persons who help with samples collecting: Bingkuan Liang, Canwei Xia, Chenxi Jia, Chunfa Zhou, Dai Cai, Fashen Zou, Guodong Wang, He Bu, Jianqiang Li, Jianrong Guo, Jiansheng Lin, Jiliang Xu, Jie Xu, Jong-Ryol Chong, Jun Xu, Kun Wu, Lu Huang, Ming Ma, Mingfu Li, Noritaka Ichida, Shicheng Lv, Songlin Huang, Tao He, Tianlin Zhou, Wei Liang, Weishi Liu, Wenbo Liao, Wenhong Deng, Xiangjiang Zhan, Xianglin Liu, Xiaodong Li, Ying Liu, Yinghong Hao, Yiqiang Fu, Yongqiang Zhang, Yumin Guo. We are very grateful to Dr. Nigel Collar for his valuable comments on an early manuscript of this work.

## DATA ACCESSIBILITY

This information will become available upon the acceptance of the manuscript.

Sequences deposited at GenBank: accession number xxxxx-xxxxx

Phenotype, distribution, stable isotope and microsatellite genotypes available at: Dryad Doi: xxx

