## Supplementary material for "Regionally divergent drivers of historical diversification in the late Quaternary in a widely distributed generalist species, the common pheasant *Phasianus colchicus*": See Supporting information for laboratory protocol and sequence editing.

***Correspondence**

**Laboratory protocols**

We extracted gross genomic DNA using TIANamp Genomic DNA kit (TIANGEN, China), following the standard extraction protocol. We measured DNA concentration and purity using a nucleic acid and protein analyzer (NanoDrop2000, ThermoFisher). We sequenced the two mtDNA loci cytochrome *b* (*cyt b*), and control region (*CR*) (Randi *et al.*, 2000; Kimball *et al.*, 2009), and additionally seven nuclear introns. These nuclear loci were widely used for Galliformes (Armstrong *et al.*, 2001; Kimball *et al.*, 2009) and included Aldolase b intron 6 (*AldB*), Beta-fibrinogen Intron 4 (*Bfibex*), Clathrin heavy polypeptide intron 6 (*CLTC*), Dimerization cofactor of HNF1 intron 3 (*DCoH*), High mobility group 17 intron 2 (*HMG*), Ovomucoid intron G (*OvoG*) and Serpin peptidase inhibitor clade C intron 5 (*SerpinC*) (Table S2). A touchdown PCR protocol was performed on a Veriti96 PCR thermal cycler system (ABI, USA) using 10 μl reaction volumes (KOD, China) (Table S3). The resulting PCR products were sequenced from both ends using the same primers as for PCR on an ABI3730XL sequencer (Applied Biosystems, USA) by MajorBio, P.R. China.

**Sequence alignments and parameter estimation**

We assembled forward and reverse amplicons and detected heterozygotes in nuclear genes using SeqMan v7.1.0.44 (Swindell and Plasterer, 1997). Alleles for nuclear introns were inferred with DnaSP v5.10.01 (Librado and Rozas, 2009). Sequences alignment was done in MEGA6 (Tamura et al., 2013) and partitioned them into three datasets for subsequent analyses: (1) mtDNA, (2) nuclear genes and (3)concatenated dataset (mtDNA+nuclear genes). Standard genetic diversity indices were estimated with DnaSP. Best-fitting models of nucleotide substitution were evaluated for each locus based on Akaike Information Criterion (AIC) using jModelTest v2.1.6 (Darriba et al., 2012).

**Phylogeographic analysis**

To visualize genetic variation, we constructed a median‐joining haplotype networks for each locus separately in PopART v4.8.4 (Leigh & Bryant, 2015). Furthermore, we generated a neighbor-net phylogenetic network based on all nuclear loci in SplitsTree v4.14.4 (Huson & Bryant, 2006).

A dated gene tree based on the mtDNA dataset was reconstructed in BEAST v2.4.7 (Bouckaert *et al.*, 2014) using linked tree models for the two mtDNA loci, but unlinked site models and clock rates. The approach of Li et al. (2010) was followed to estimate substitution rates of the different loci. We firstly computed the overall mean of genetic distances of each locus among all samples including outgroup in MEGA with 1000 bootstrap replicates. Then the ratio of genetic distance between each locus and *cyt b* was calculated. Finally, we multiplied the resulting ratio by average divergence rate of *cyt b* for Galliformes 0.0238 substitutions/site/million year (Weir & Schluter, 2008) to estimate substitution rates for each locus. Inferred best-fitting models of nucleotide evolution for each locus were used as site models. A relaxed uncorrelated lognormal distribution was selected for the clock model. A log normal distribution with standard deviation of 0.05 was used as the prior distribution of the clock rates with the following mean values (in real space) of 0.0119 substitution/site/Million years for *cyt* *b* and 0.0315 substitution/site/Million years for *CR* (Tab. S5). MCMC analyses were performed three times independently using a coalescent exponential tree model with 50 million iterations storing every 1,000 generations.

A dated species tree for the concatenated dataset was reconstructed using *BEAST v2.4.7 (Bouckaert *et al.*, 2014) based on the two mtDNA loci and the alleles of six nuclear introns. *AldB* located on the Z chromosome was excluded from this analyses as information on sex was not available for all individuals. Tree models were linked for the two mtDNA loci but unlinked for the nuclear introns, while site models and clock rates were unlinked for all loci. Based on the results of the mtDNA gene tree estimated with BEAST (see below), samples of eight different evolutionary lineages were set as ‘taxa’. Best-fitting models of nucleotide evolution were implemented as site models for each locus (Tab. S5). A relaxed lognormal distribution was selected and substitution rates of each locus were implemented as a mean value (in real space) of a log normal distribution with a standard deviation of 0.05 with locus specific mean substitution rates (Table S5). We selected Yule model for the species tree prior and a linear with constant root prior for the population function. Four independent runs were run with 1 billion generations stored every 0.2 million generations.

*BEAST was moreover used to reconstruct the colonization routes of the eight evolutionary lineages based on the concatenated dataset as above in dated species tree analyses. Coordinates representing the central locality of each lineage were implemented as a trait using GEO_SPhERE and BEAST-CLASSIC packages (Lemey *et al.*, 2009). Three independent chains were run for two billion generations and sampled every 0.1 million generation. We used the combined MCC tree (see below) with spherical geography in SPREAD v1.0.7 (Bielejec *et al.*, 2011) to infer the colonization routes of common pheasants, which were displayed in google-earth v2 (Yamagishi *et al.*, 2010).

For both BEAST and *BEAST analyses, Tracer v1.6 (Rambaut *et al.*, 2013) was used to check for adequate effective sample sizes (ESS) of the posterior distribution of each parameter and to assess convergence among the independent runs. The runs were the combined using LogCombiner v2.4.7 (Bouckaert *et al.*, 2014) with 10% each discarded as burn-in. The combined posterior distributions were summarized to produce a maximum clade credibility (MCC) tree in TreeAnnotator v1.8.2 (Drummond & Rambaut, 2007) using 10,000 randomly chosen trees in the BEAST analyses. Analyses were performed on the CIPRES Science Gateway (Miller *et al.*, 2010) and the resulting trees were visualized in FigTree v1.4 (Rambaut, 2008) and DensiTree v2.0 (Bouckaert & Heled, 2014) .

**Demographic and migration analyses**

Changes in effective population size (*Ne*) through time in the different evolutionary lineages (see below) were reconstructed with extended Bayesian Skyline Plot (EBSP) in BEAST v2.4.7. using the concatenated dataset. Due to low sample sizes, no inference of demographic history was performed for the *tarimensis* and the *formosanus* groups. Again, tree models were linked for the two mtDNA loci but unlinked for the nuclear introns, while site models and clock rates were unlinked for all loci. The same priors as were implemented above for substitution models and clock rates. A coalescent extend Bayesian Skyline tree prior was used. The MCMC analysis of each group was run for 100 million generations with sampling each 1000 generations.

The parameter sum (indicators.alltrees) was check to infer for the most likely numbers of demographic changes. Convergence and ESS value were assessed again in Tracer v1.6. Skyline plots were generated using R scripts available at: <https://github.com/CompEvol/beast2/tree/master/doc/tutorials/EBSP>. Since EBSP analyses assume no gene flow after divergence between lineages, we additionally inferred non-equilibrium scenarios between parapatric lineages based on an isolation-with-migration (IM) model. This model allowed us to assess migration rates, effective population size and divergence time. We run IM analyses on each pair of parapatric lineages populations using IMa2 (Hey, 2010a, b). Population string was defined based on the results of the phylogeographic analyses (see below). The substitution rates (per locus per year) and their interval for each locus as estimated using genetic distance described above in ‘Phylogeographic analysis’ and their standard deviation were used as mutation rates and their ranges. A HKY model was set as substitution model. Generation time was set to 2 as sexual maturity is reached after two years in common pheasants. We run multiple simulations to test for appropriate upper bounds of migration rates, population sizes and time of population splits based on the results of the posterior distribution. Each analysis was executed for 10 million generations, storing every 1,000 MCMC steps.

**Species delimitation**

We moreover conducted a species delimitation analyses in BPP v.3.4 (Yang, 2015). We used the A10 model (speciesdelimitation=1, speciestree=0) for species delimitation using the species-tree of the *BEAST analyses (see below) as the user-specified guide tree (Yang & Rannala, 2010; Rannala & Yang, 2013) treating the eight distinct evolutionary lineages within common pheasant as potential species with equal prior probabilities to all potential species delimitation models. We tested three different sets of gamma priors for the population size (θ) and divergence time (τ) parameters: Γ( 1, 1000) for both parameters, θ~Γ(1, 10) and τ~Γ (1, 1000), as well as θ~Γ (1, 1000) and τ~Γ (1, 10). Substitution rates as estimate above were used as locus rates with both mtDNA loci treated together as a single recombination unit. Three independent runs were performed 100,000 generations with sampling every 10 samples, discarding the first 10,000 samples as burn-in.

**Table S1.** List of all samples used in this study including sample number and information on origin and locality, sample type, sex as well as GenBank accession numbers. Blood and muscle samples were stored in ethanol, while feathers were keep in dry paper envelopes, and then kept at -40^o^C at the Museum of Biology of Sun Yat-sen University, Guangzhou, P.R. China, for long-term storage. GenBank accession number for each sequences will be available when this manuscript is accepted.

| **Number** | **Subspecies** | **abbr.** | **Location** | **Longi-tude** | **Lati-tude** | **Sample Type** | **Sex** | ***CR*** | ***Cyt b*** | ***AldB*** | ***Bfibex*** | ***CLTC*** | ***DCoH*** | ***HMG17*** | ***SerpinC*** | ***OvoG*** |
| --- | --- | --- | --- | --- | --- | --- | --- | --- | --- | --- | --- | --- | --- | --- | --- | --- |
| SYSb002945 | *alaschanicus* | ALA | Alxa Left Banner, Inner Mongolia, China | 111.17 | 40.70 | Muscle |  |  |  |  |  |  |  |  |  |  |
| SYSb002946 | *alaschanicus* | ALA | Alxa Left Banner, Inner Mongolia, China | 111.17 | 40.70 | Muscle |  |  |  |  |  |  |  |  |  |  |
| SYSb002947 | *alaschanicus* | ALA | Alxa Left Banner, Inner Mongolia, China | 111.17 | 40.70 | Muscle |  |  |  |  |  |  |  |  |  |  |
| SYSb001733 | *bianchii* | BIA | Captive population |  |  | Feather | ♀ |  |  |  |  |  |  |  |  |  |
| SYSb001735 | *bianchii* | BIA | Captive population |  |  | Feather | ♀ |  |  |  |  |  |  |  |  |  |
| SYSb001737 | *bianchii* | BIA | Captive population |  |  | Feather | ♀ |  |  |  |  |  |  |  |  |  |
| SYSb001738 | *bianchii* | BIA | Captive population |  |  | Feather | ♀ |  |  |  |  |  |  |  |  |  |
| SYSb001739 | *bianchii* | BIA | Captive population |  |  | Feather | ♀ |  |  |  |  |  |  |  |  |  |
| SYSb001751 | *bianchii* | BIA | Captive population |  |  | Feather | ♂ |  |  |  |  |  |  |  |  |  |
| SYSb001771 | *bianchii* | BIA | Captive population |  |  | Feather | ♀ |  |  |  |  |  |  |  |  |  |
| SYSb001777 | *bianchii* | BIA | Dusambe Zoo,Tajikistan | 68.78 | 38.54 | Feather | ♀ |  |  |  |  |  |  |  |  |  |
| SYSb001785 | *bianchii* | BIA | Dusambe Zoo,Tajikistan | 68.78 | 38.54 | Feather | ♂ |  |  |  |  |  |  |  |  |  |
| SYSb001789 | *bianchii* | BIA | Dusambe Zoo,Tajikistan | 68.78 | 38.54 | Feather | ♂ |  |  |  |  |  |  |  |  |  |
| SYSb005765 | *colchicus* | COL | Barda, Azerbaijan | 47.14 | 40.36 | DNA |  |  |  |  |  |  |  |  |  |  |
| SYSb005766 | *colchicus* | COL | Barda, Azerbaijan | 47.14 | 40.36 | DNA |  |  |  |  |  |  |  |  |  |  |
| SYSb005767 | *colchicus* | COL | Barda, Azerbaijan | 47.14 | 40.36 | DNA |  |  |  |  |  |  |  |  |  |  |
| SYSb005768 | *colchicus* | COL | Barda, Azerbaijan | 47.14 | 40.36 | DNA |  |  |  |  |  |  |  |  |  |  |
| SYSb005769 | *colchicus* | COL | Barda, Azerbaijan | 47.14 | 40.36 | DNA |  |  |  |  |  |  |  |  |  |  |
| SYSb005770 | *colchicus* | COL | Barda, Azerbaijan | 47.14 | 40.36 | DNA |  |  |  |  |  |  |  |  |  |  |
| SYSb005771 | *colchicus* | COL | Barda, Azerbaijan | 47.14 | 40.36 | DNA |  |  |  |  |  |  |  |  |  |  |
| SYSb005772 | *colchicus* | COL | Barda, Azerbaijan | 47.14 | 40.36 | DNA |  |  |  |  |  |  |  |  |  |  |
| SYSb004573 | *decollatus* | DEC | Sandu, Guizhou, China | 108.02 | 25.61 | Muscle |  |  |  |  |  |  |  |  |  |  |
| SYSb004964 | *decollatus* | DEC | Sandu, Guizhou, China | 108.02 | 25.61 | Muscle | ♂ |  |  |  |  |  |  |  |  |  |
| SYSb004426 | *decollatus* | DEC | Shiqian, Guizhou, China | 108.22 | 27.51 | Blood | ♂ |  |  |  |  |  |  |  |  |  |
| SYSb004437 | *decollatus* | DEC | Shiqian, Guizhou, China | 108.22 | 27.51 | Blood |  |  |  |  |  |  |  |  |  |  |
| SYSb004424 | *decollatus* | DEC | Zhenyuan, Guizhou, China | 108.43 | 27.05 | Blood | ♂ |  |  |  |  |  |  |  |  |  |
| SYSb004425 | *decollatus* | DEC | Zhenyuan, Guizhou, China | 108.43 | 27.05 | Blood | ♀ |  |  |  |  |  |  |  |  |  |
| SYSb004954 | *decollatus* | DEC | Zhenyuan, Guizhou, China | 108.43 | 27.05 | Blood |  |  |  |  |  |  |  |  |  |  |
| SYSb005773 | *elegans* | ELE | Dali, Yunnan, China | 100.27 | 25.59 | DNA |  |  |  |  |  |  |  |  |  |  |
| SYSb005680 | *elegans* | ELE | Kunming, Yunnan, China | 102.83 | 24.88 | DNA |  |  |  |  |  |  |  |  |  |  |
| SYSb004896 | *elegans* | ELE | Leshan, Sichuan, China | 103.77 | 29.56 | Muscle | ♂ |  |  |  |  |  |  |  |  |  |
| SYSb004639 | *elegans* | ELE | Nanjian, Yunan, China | 100.51 | 25.04 | Blood | ♀ |  |  |  |  |  |  |  |  |  |
| SYSb004640 | *elegans* | ELE | Nanjian, Yunan, China | 100.51 | 25.04 | Blood | ♀ |  |  |  |  |  |  |  |  |  |
| SYSb004739 | *elegans* | ELE | Shangri-La, Yunnan, China | 99.70 | 27.83 | Muscle |  |  |  |  |  |  |  |  |  |  |
| SYSb004740 | *elegans* | ELE | Shangri-La, Yunnan, China | 99.70 | 27.83 | Muscle |  |  |  |  |  |  |  |  |  |  |
| SYSb004642 | *elegans* | ELE | Xinping, Yunnan, China | 103.23 | 22.78 | Blood |  |  |  |  |  |  |  |  |  |  |
| SYSb004643 | *elegans* | ELE | Xinping, Yunnan, China | 103.23 | 22.78 | Blood |  |  |  |  |  |  |  |  |  |  |
| SYSb005040 | *elegans* | ELE | Zhaotong, Yunnan, China | 103.72 | 27.34 | Muscle |  |  |  |  |  |  |  |  |  |  |
| SYSb005072 | *elegans* | ELE | Zhaotong, Yunnan, China | 103.72 | 27.34 | Muscle |  |  |  |  |  |  |  |  |  |  |
| SYSb002229 | *formosanus* | FOR | Taiwan, China | 121.08 | 23.90 | Muscle |  |  |  |  |  |  |  |  |  |  |
| SYSb002230 | *formosanus* | FOR | Taiwan, China | 121.08 | 23.90 | Muscle |  |  |  |  |  |  |  |  |  |  |
| SYSb002233 | *formosanus* | FOR | Taiwan, China | 121.08 | 23.90 | Muscle |  |  |  |  |  |  |  |  |  |  |
| SYSb002235 | *formosanus* | FOR | Taiwan, China | 121.08 | 23.90 | Muscle |  |  |  |  |  |  |  |  |  |  |
| SYSb002242 | *formosanus* | FOR | Taiwan, China | 121.08 | 23.90 | Muscle |  |  |  |  |  |  |  |  |  |  |
| SYSb002244 | *formosanus* | FOR | Taiwan, China | 121.08 | 23.90 | Muscle |  |  |  |  |  |  |  |  |  |  |
| SYSb002246 | *formosanus* | FOR | Taiwan, China | 121.08 | 23.90 | Muscle |  |  |  |  |  |  |  |  |  |  |
| SYSb002247 | *formosanus* | FOR | Taiwan, China | 121.08 | 23.90 | Muscle |  |  |  |  |  |  |  |  |  |  |
| SYSb001044 | *hagenbecki* | HAG | Gobi, Mongolia | 109.19 | 43.80 | Feather |  |  |  |  |  |  |  |  |  |  |
| **Number** | **Subspecies** | **abbr.** | **Location** | **Longi-tude** | **Lati-tude** | **Sample Type** | **Gen-der** | ***CR*** | ***Cyt b*** | ***AldB*** | ***Bfibex*** | ***CLTC*** | ***DCoH*** | ***HMG17*** | ***SerpinC*** | ***OvoG*** |
| SYSb001045 | *hagenbecki* | HAG | Gobi, Mongolia | 109.19 | 43.80 | Feather |  |  |  |  |  |  |  |  |  |  |
| SYSb001048 | *hagenbecki* | HAG | Gobi, Mongolia | 109.19 | 43.80 | Feather |  |  |  |  |  |  |  |  |  |  |
| SYSb001049 | *hagenbecki* | HAG | Gobi, Mongolia | 109.19 | 43.80 | Feather |  |  |  |  |  |  |  |  |  |  |
| SYSb001050 | *hagenbecki* | HAG | Gobi, Mongolia | 109.19 | 43.80 | Feather |  |  |  |  |  |  |  |  |  |  |
| SYSb001052 | *hagenbecki* | HAG | Gobi, Mongolia | 109.19 | 43.80 | Feather |  |  |  |  |  |  |  |  |  |  |
| SYSb001055 | *hagenbecki* | HAG | Gobi, Mongolia | 109.19 | 43.80 | Feather |  |  |  |  |  |  |  |  |  |  |
| SYSb004749 | *karpowi* | KAR | Chengde, Hebei, China | 117.74 | 41.90 | Muscle |  |  |  |  |  |  |  |  |  |  |
| SYSb004875 | *karpowi* | KAR | Chengde, Hebei, China | 117.74 | 41.90 | Muscle |  |  |  |  |  |  |  |  |  |  |
| SYSb004971 | *karpowi* | KAR | Chengde, Hebei, China | 117.74 | 41.90 | Muscle |  |  |  |  |  |  |  |  |  |  |
| SYSb004478 | *karpowi* | KAR | Chifeng, Inner Mongolia, China | 118.65 | 44.25 | Muscle | ♂ |  |  |  |  |  |  |  |  |  |
| SYSb004479 | *karpowi* | KAR | Chifeng, Inner Mongolia, China | 118.65 | 44.25 | Muscle | ♀ |  |  |  |  |  |  |  |  |  |
| SYSb005018 | *karpowi* | KAR | Dandong, Liaoning, China | 124.07 | 40.45 | Muscle | ♂ |  |  |  |  |  |  |  |  |  |
| SYSb005774 | *karpowi* | KAR | Huludao, Liaoning, China | 120.76 | 40.64 | DNA |  |  |  |  |  |  |  |  |  |  |
| SYSb005002 | *kiansuensis* | KIA | Huhehot, Inner Mongolia, China | 111.75 | 40.84 | Muscle | ♂ |  |  |  |  |  |  |  |  |  |
| SYSb004509 | *kiansuensis* | KIA | Jiaocheng, Shanxi, China | 111.36 | 37.90 | Muscle |  |  |  |  |  |  |  |  |  |  |
| SYSb004511 | *kiansuensis* | KIA | Jiaocheng, Shanxi, China | 111.36 | 37.90 | Muscle |  |  |  |  |  |  |  |  |  |  |
| SYSb004512 | *kiansuensis* | KIA | Jiaocheng, Shanxi, China | 111.36 | 37.90 | Muscle | ♂ |  |  |  |  |  |  |  |  |  |
| SYSb004513 | *kiansuensis* | KIA | Jiaocheng, Shanxi, China | 111.36 | 37.90 | Muscle | ♂ |  |  |  |  |  |  |  |  |  |
| SYSb004514 | *kiansuensis* | KIA | Jiaocheng, Shanxi, China | 111.36 | 37.90 | Muscle |  |  |  |  |  |  |  |  |  |  |
| SYSb004537 | *kiansuensis* | KIA | Jiaocheng, Shanxi, China | 111.36 | 37.90 | Muscle | ♂ |  |  |  |  |  |  |  |  |  |
| SYSb004422 | *kiansuensis* | KIA | Luya Mountain, Shanxi, China | 111.93 | 38.73 | Muscle | ♂ |  |  |  |  |  |  |  |  |  |
| SYSb004378 | *kiansuensis* | KIA | Ulanqab, Inner Mongolia, China | 112.51 | 40.51 | Muscle | ♀ |  |  |  |  |  |  |  |  |  |
| SYSb004551 | *kiansuensis* | KIA | Zhulu, Hebei, China | 115.21 | 40.38 | Muscle |  |  |  |  |  |  |  |  |  |  |
| SYSb004625 | *kiansuensis* | KIA | Zhulu, Hebei, China | 115.21 | 40.38 | Muscle | ♂ |  |  |  |  |  |  |  |  |  |
| SYSb005776 | *mongolicus* | MON | Huocheng, Xinjiang, China | 80.88 | 44.06 | DNA |  |  |  |  |  |  |  |  |  |  |
| SYSb006581 | *mongolicus* | MON | Huocheng, Xinjiang, China | 80.88 | 44.06 | Muscle |  |  |  |  |  |  |  |  |  |  |
| SYSb006582 | *mongolicus* | MON | Huocheng, Xinjiang, China | 80.88 | 44.06 | Muscle |  |  |  |  |  |  |  |  |  |  |
| SYSb006583 | *mongolicus* | MON | Huocheng, Xinjiang, China | 80.88 | 44.06 | Muscle |  |  |  |  |  |  |  |  |  |  |
| SYSb006584 | *mongolicus* | MON | Huocheng, Xinjiang, China | 80.88 | 44.06 | Muscle |  |  |  |  |  |  |  |  |  |  |
| SYSb006585 | *mongolicus* | MON | Huocheng, Xinjiang, China | 80.88 | 44.06 | Muscle |  |  |  |  |  |  |  |  |  |  |
| SYSb006587 | *mongolicus* | MON | Huocheng, Xinjiang, China | 80.88 | 44.06 | Muscle |  |  |  |  |  |  |  |  |  |  |
| SYSb006588 | *mongolicus* | MON | Huocheng, Xinjiang, China | 80.88 | 44.06 | Muscle |  |  |  |  |  |  |  |  |  |  |
| SYSb006589 | *mongolicus* | MON | Huocheng, Xinjiang, China | 80.88 | 44.06 | Muscle |  |  |  |  |  |  |  |  |  |  |
| SYSb006590 | *mongolicus* | MON | Huocheng, Xinjiang, China | 80.88 | 44.06 | Muscle |  |  |  |  |  |  |  |  |  |  |
| SYSb006591 | *mongolicus* | MON | Huocheng, Xinjiang, China | 80.88 | 44.06 | Muscle |  |  |  |  |  |  |  |  |  |  |
| SYSb006592 | *mongolicus* | MON | Huocheng, Xinjiang, China | 80.88 | 44.06 | Muscle |  |  |  |  |  |  |  |  |  |  |
| SYSb006593 | *mongolicus* | MON | Huocheng, Xinjiang, China | 80.88 | 44.06 | Muscle |  |  |  |  |  |  |  |  |  |  |
| SYSb006594 | *mongolicus* | MON | Huocheng, Xinjiang, China | 80.88 | 44.06 | Muscle |  |  |  |  |  |  |  |  |  |  |
| SYSb006595 | *mongolicus* | MON | Huocheng, Xinjiang, China | 80.88 | 44.06 | Muscle |  |  |  |  |  |  |  |  |  |  |
| SYSb006596 | *mongolicus* | MON | Huocheng, Xinjiang, China | 80.88 | 44.06 | Muscle |  |  |  |  |  |  |  |  |  |  |
| SYSb005777 | *mongolicus* | MON | Wenquan, Xinjiang, China | 81.04 | 44.96 | DNA |  |  |  |  |  |  |  |  |  |  |
| SYSb005778 | *mongolicus* | MON | Wenquan, Xinjiang, China | 81.04 | 44.96 | DNA |  |  |  |  |  |  |  |  |  |  |
| SYSb004719 | *pallasi* | PAL | Daan, Jilin, China | 124.25 | 45.47 | Muscle | ♀ |  |  |  |  |  |  |  |  |  |
| SYSb004777 | *pallasi* | PAL | Daqing, Heilongjiang, China | 124.76 | 46.89 | Muscle |  |  |  |  |  |  |  |  |  |  |
| SYSb004722 | *pallasi* | PAL | Dehui, Jilin, China | 125.73 | 44.50 | Muscle |  |  |  |  |  |  |  |  |  |  |
| SYSb004419 | *pallasi* | PAL | Jiamusi, Heilongjiang, China | 132.03 | 47.22 | Muscle |  |  |  |  |  |  |  |  |  |  |
| SYSb005779 | *pallasi* | PAL | Xianghai, Jilin, China | 122.56 | 44.91 | Muscle |  |  |  |  |  |  |  |  |  |  |
| SYSb005035 | *pallasi* | PAL | Xingkai Lake, Heilongjiang, China | 132.38 | 45.00 | Muscle | ♂ |  |  |  |  |  |  |  |  |  |
| SYSb004617 | *pallasi* | PAL | Xunke, Heilongjiang, China | 128.48 | 49.56 | Muscle |  |  |  |  |  |  |  |  |  |  |
| SYSb004503 | *pallasi* | PAL | Yakeshi, Inner Mongolia, China | 120.71 | 49.29 | Muscle |  |  |  |  |  |  |  |  |  |  |
| SYSb004947 | *pallasi* | PAL | Yakeshi, Inner Mongolia, China | 120.71 | 49.29 | Muscle |  |  |  |  |  |  |  |  |  |  |
| SYSb004720 | *pallasi* | PAL | Yanji, Jilin, China | 129.51 | 42.87 | Muscle | ♂ |  |  |  |  |  |  |  |  |  |
| SYSb005781 | *persicus* | PER | Bandpey-e-sharghi, Iran | 51.35 | 35.70 | DNA |  |  |  |  |  |  |  |  |  |  |
| SYSb005783 | *persicus* | PER | Chaboksar, Iran | 50.56 | 36.97 | DNA |  |  |  |  |  |  |  |  |  |  |
| SYSb005784 | *persicus* | PER | Chaboksar, Iran | 50.56 | 36.97 | DNA |  |  |  |  |  |  |  |  |  |  |
| SYSb006570 | *persicus* | PER | Mazandaran, Iran | 53.05 | 36.56 | Blood |  |  |  |  |  |  |  |  |  |  |
| SYSb006574 | *persicus* | PER | Mazandaran, Iran | 53.05 | 36.56 | Blood |  |  |  |  |  |  |  |  |  |  |
| SYSb006575 | *persicus* | PER | Mazandaran, Iran | 53.05 | 36.56 | Blood |  |  |  |  |  |  |  |  |  |  |
| SYSb005785 | *persicus* | PER | Minoo Dasht, Iran | 55.37 | 37.21 | DNA |  |  |  |  |  |  |  |  |  |  |
| SYSb005786 | *persicus* | PER | Minoo Dasht, Iran | 55.37 | 37.21 | DNA |  |  |  |  |  |  |  |  |  |  |
| SYSb005787 | *persicus* | PER | Minoo Dasht, Iran | 55.37 | 37.21 | DNA |  |  |  |  |  |  |  |  |  |  |
| SYSb005780 | *persicus* | PER | Ramsar, Iran | 50.64 | 36.93 | DNA |  |  |  |  |  |  |  |  |  |  |
| SYSb005782 | *persicus* | PER | Rasht, Iran | 49.60 | 37.31 | DNA |  |  |  |  |  |  |  |  |  |  |
| SYSb005788 | *principalis* | PRI | Dargaz, Iran | 59.11 | 37.44 | DNA |  |  |  |  |  |  |  |  |  |  |
| SYSb005789 | *principalis* | PRI | Dargaz, Iran | 59.11 | 37.44 | DNA |  |  |  |  |  |  |  |  |  |  |
| SYSb005790 | *principalis* | PRI | Sarakhs, Iran | 61.15 | 36.54 | DNA |  |  |  |  |  |  |  |  |  |  |
| SYSb005791 | *principalis* | PRI | Sarakhs, Iran | 61.15 | 36.54 | DNA |  |  |  |  |  |  |  |  |  |  |
| SYSb005792 | *principalis* | PRI | Sarakhs, Iran | 61.15 | 36.54 | DNA |  |  |  |  |  |  |  |  |  |  |
| SYSb005793 | *principalis* | PRI | Sarakhs, Iran | 61.15 | 36.54 | DNA |  |  |  |  |  |  |  |  |  |  |
| SYSb005794 | *principalis* | PRI | Sarakhs, Iran | 61.15 | 36.54 | DNA |  |  |  |  |  |  |  |  |  |  |
| SYSb005795 | *principalis* | PRI | Sarakhs, Iran | 61.15 | 36.54 | DNA |  |  |  |  |  |  |  |  |  |  |
| SYSb005796 | *principalis* | PRI | Sarakhs, Iran | 61.15 | 36.54 | DNA |  |  |  |  |  |  |  |  |  |  |
| SYSb004452 | *satscheuensis* | SAT | Yumen, Gansu, China | 97.05 | 40.29 | Muscle |  |  |  |  |  |  |  |  |  |  |
| SYSb004453 | *satscheuensis* | SAT | Yumen, Gansu, China | 97.05 | 40.29 | Muscle | ♂ |  |  |  |  |  |  |  |  |  |
| SYSb004454 | *satscheuensis* | SAT | Yumen, Gansu, China | 97.05 | 40.29 | Muscle | ♂ |  |  |  |  |  |  |  |  |  |
| SYSb004878 | *satscheuensis* | SAT | Yumen, Gansu, China | 97.05 | 40.29 | Muscle |  |  |  |  |  |  |  |  |  |  |
| SYSb004893 | *satscheuensis* | SAT | Yumen, Gansu, China | 97.05 | 40.29 | Muscle |  |  |  |  |  |  |  |  |  |  |
| SYSb004894 | *satscheuensis* | SAT | Yumen, Gansu, China | 97.05 | 40.29 | Feather | ♂ |  |  |  |  |  |  |  |  |  |
| SYSb004895 | *satscheuensis* | SAT | Yumen, Gansu, China | 97.05 | 40.29 | Muscle | ♀ |  |  |  |  |  |  |  |  |  |
| SYSb005056 | *satscheuensis* | SAT | Yumen, Gansu, China | 97.05 | 40.29 | Muscle | ♀ |  |  |  |  |  |  |  |  |  |
| SYSb005797 | *satscheuensis* | SAT | Yumen, Gansu, China | 97.05 | 40.29 | DNA |  |  |  |  |  |  |  |  |  |  |
| SYSb000336 | *shawi* | SHA | Aksu, Xinjiang, China | 80.26 | 41.17 | Muscle | ♂ |  |  |  |  |  |  |  |  |  |
| SYSb000337 | *shawi* | SHA | Aksu, Xinjiang, China | 80.26 | 41.17 | Muscle | ♂ |  |  |  |  |  |  |  |  |  |
| SYSb000338 | *shawi* | SHA | Aksu, Xinjiang, China | 80.26 | 41.17 | Muscle | ♂ |  |  |  |  |  |  |  |  |  |
| SYSb004555 | *shawi* | SHA | Aksu, Xinjiang, China | 80.26 | 41.17 | Muscle | ♂ |  |  |  |  |  |  |  |  |  |
| SYSb004558 | *shawi* | SHA | Aksu, Xinjiang, China | 80.26 | 41.17 | Muscle | ♂ |  |  |  |  |  |  |  |  |  |
| SYSb004955 | *shawi* | SHA | Aksu, Xinjiang, China | 80.26 | 41.17 | Muscle | ♂ |  |  |  |  |  |  |  |  |  |
| SYSb005166 | *shawi* | SHA | Aksu, Xinjiang, China | 80.26 | 41.17 | DNA |  |  |  |  |  |  |  |  |  |  |
| SYSb005167 | *shawi* | SHA | Aksu, Xinjiang, China | 80.26 | 41.17 | DNA |  |  |  |  |  |  |  |  |  |  |
| SYSb005168 | *shawi* | SHA | Aksu, Xinjiang, China | 80.26 | 41.17 | DNA |  |  |  |  |  |  |  |  |  |  |
| SYSb005169 | *shawi* | SHA | Aksu, Xinjiang, China | 80.26 | 41.17 | DNA |  |  |  |  |  |  |  |  |  |  |
| SYSb005170 | *shawi* | SHA | Aksu, Xinjiang, China | 80.26 | 41.17 | DNA |  |  |  |  |  |  |  |  |  |  |
| SYSb005798 | *shawi* | SHA | Aksu, Xinjiang, China | 80.26 | 41.17 | DNA |  |  |  |  |  |  |  |  |  |  |
| SYSb005799 | *shawi* | SHA | Aksu, Xinjiang, China | 80.26 | 41.17 | DNA |  |  |  |  |  |  |  |  |  |  |
| SYSb005171 | *shawi* | SHA | Kashgar, Xinjiang, China | 75.99 | 39.46 | DNA |  |  |  |  |  |  |  |  |  |  |
| SYSb005172 | *shawi* | SHA | Kashgar, Xinjiang, China | 75.99 | 39.46 | DNA |  |  |  |  |  |  |  |  |  |  |
| SYSb004542 | *shawi* | SHA | Yanqi Basin, Xinjiang, China | 86.44 | 41.93 | Muscle |  |  |  |  |  |  |  |  |  |  |
| SYSb004494 | *strauchi* | STR | Baoji, Shaanxi, China | 107.13 | 34.64 | Muscle |  |  |  |  |  |  |  |  |  |  |
| SYSb004974 | *strauchi* | STR | Datong, Qinghai, China | 101.69 | 36.93 | Muscle |  |  |  |  |  |  |  |  |  |  |
| SYSb004948 | *strauchi* | STR | Foping, Shaanxi, China | 107.99 | 33.52 | Feather | ♂ |  |  |  |  |  |  |  |  |  |
| SYSb004447 | *strauchi* | STR | Guangyuan, Sichuan, China | 104.76 | 32.58 | Muscle | ♂ |  |  |  |  |  |  |  |  |  |
| SYSb004448 | *strauchi* | STR | Guangyuan, Sichuan, China | 105.24 | 32.58 | Muscle | ♂ |  |  |  |  |  |  |  |  |  |
| SYSb004650 | *strauchi* | STR | Guangyuan, Sichuan, China | 104.76 | 32.58 | Muscle |  |  |  |  |  |  |  |  |  |  |
| SYSb005048 | *strauchi* | STR | Guyuan, Ningxia, China | 106.24 | 36.02 | Muscle |  |  |  |  |  |  |  |  |  |  |
| SYSb004726 | *strauchi* | STR | Huangyuan, Qinghai, China | 101.23 | 36.68 | Muscle |  |  |  |  |  |  |  |  |  |  |
| SYSb004999 | *strauchi* | STR | Huangyuan, Qinghai, China | 101.23 | 36.68 | Muscle | ♂ |  |  |  |  |  |  |  |  |  |
| SYSb004687 | *strauchi* | STR | Qingyang, Gansu, China | 107.65 | 35.72 | Muscle |  |  |  |  |  |  |  |  |  |  |
| SYSb004688 | *strauchi* | STR | Qingyang, Gansu, China | 107.65 | 35.72 | Muscle |  |  |  |  |  |  |  |  |  |  |
| SYSb005068 | *strauchi* | STR | Qingyang, Gansu, China | 107.65 | 35.72 | Muscle |  |  |  |  |  |  |  |  |  |  |
| SYSb005069 | *strauchi* | STR | Qingyang, Gansu, China | 107.65 | 35.72 | Muscle |  |  |  |  |  |  |  |  |  |  |
| SYSb004993 | *strauchi* | STR | Xining, Qinghai, China | 101.57 | 36.47 | Muscle | ♂ |  |  |  |  |  |  |  |  |  |
| SYSb005036 | *suehschanensis* | SUE | Qingchuan, Sichuan, China | 105.24 | 32.58 | Muscle | ♀ |  |  |  |  |  |  |  |  |  |
| SYSb005037 | *suehschanensis* | SUE | Qingchuan, Sichuan, China | 105.24 | 32.58 | Muscle | ♂ |  |  |  |  |  |  |  |  |  |
| SYSb004912 | *suehschanensis* | SUE | Suining, Sichuan, China | 105.59 | 30.54 | Muscle |  |  |  |  |  |  |  |  |  |  |
| SYSb004914 | *suehschanensis* | SUE | Suining, Sichuan, China | 105.59 | 30.54 | Muscle |  |  |  |  |  |  |  |  |  |  |
| SYSb004783 | *takatsukasae* | TAK | Fangcheng, Guangxi, China | 108.35 | 21.69 | Muscle |  |  |  |  |  |  |  |  |  |  |
| SYSb004784 | *takatsukasae* | TAK | Fangcheng, Guangxi, China | 108.35 | 21.69 | Muscle |  |  |  |  |  |  |  |  |  |  |
| SYSb005173 | *talischensis* | TAL | Abad, Iran | 54.55 | 37.34 | DNA |  |  |  |  |  |  |  |  |  |  |
| SYSb005174 | *talischensis* | TAL | Abad, Iran | 54.55 | 37.34 | DNA |  |  |  |  |  |  |  |  |  |  |
| SYSb005175 | *talischensis* | TAL | Abad, Iran | 54.55 | 37.34 | DNA |  |  |  |  |  |  |  |  |  |  |
| SYSb005176 | *talischensis* | TAL | Abad, Iran | 54.55 | 37.34 | DNA |  |  |  |  |  |  |  |  |  |  |
| SYSb005177 | *talischensis* | TAL | Abad, Iran | 54.55 | 37.34 | DNA |  |  |  |  |  |  |  |  |  |  |
| SYSb005178 | *talischensis* | TAL | Abad, Iran | 54.55 | 37.34 | DNA |  |  |  |  |  |  |  |  |  |  |
| SYSb005179 | *talischensis* | TAL | Abad, Iran | 54.55 | 37.34 | DNA |  |  |  |  |  |  |  |  |  |  |
| SYSb006569 | *talischensis* | TAL | Guilan, Iran | 49.61 | 37.32 | Blood |  |  |  |  |  |  |  |  |  |  |
| SYSb006571 | *talischensis* | TAL | Guilan, Iran | 49.61 | 37.32 | Blood |  |  |  |  |  |  |  |  |  |  |
| SYSb006572 | *talischensis* | TAL | Guilan, Iran | 49.61 | 37.32 | Blood |  |  |  |  |  |  |  |  |  |  |
| SYSb006573 | *talischensis* | TAL | Guilan, Iran | 49.61 | 37.32 | Blood |  |  |  |  |  |  |  |  |  |  |
| SYSb006577 | *talischensis* | TAL | Guilan-eshkevarat, Iran | 49.61 | 37.32 | Blood |  |  |  |  |  |  |  |  |  |  |
| SYSb006578 | *talischensis* | TAL | Guilan-eshkevarat, Iran | 49.61 | 37.32 | Blood |  |  |  |  |  |  |  |  |  |  |
| SYSb006580 | *talischensis* | TAL | Guilan-eshkevarat, Iran | 49.61 | 37.32 | Blood |  |  |  |  |  |  |  |  |  |  |
| SYSb004544 | *tarimensis* | TAR | Heshuo, Xinjiang, China | 86.86 | 42.27 | Muscle |  |  |  |  |  |  |  |  |  |  |
| SYSb004545 | *tarimensis* | TAR | Heshuo, Xinjiang, China | 86.86 | 42.27 | Muscle |  |  |  |  |  |  |  |  |  |  |
| SYSb004559 | *tarimensis* | TAR | Heshuo, Xinjiang, China | 86.86 | 42.27 | Muscle | ♂ |  |  |  |  |  |  |  |  |  |
| SYSb004543 | *tarimensis* | TAR | Weili, Xinjiang, China | 86.26 | 41.34 | Muscle |  |  |  |  |  |  |  |  |  |  |
| SYSb004724 | *torquatus* | TOR | Cangshan Mountain, Shandong, China | 121.02 | 36.93 | Muscle |  |  |  |  |  |  |  |  |  |  |
| SYSb004402 | *torquatus* | TOR | Fujian, China | 119.29 | 26.11 | Muscle |  |  |  |  |  |  |  |  |  |  |
| SYSb004390 | *torquatus* | TOR | Guangshui, Hubei, China | 113.93 | 31.84 | Muscle |  |  |  |  |  |  |  |  |  |  |
| SYSb004709 | *torquatus* | TOR | Guilin, Guangxi, China | 110.21 | 25.24 | Muscle | ♀ |  |  |  |  |  |  |  |  |  |
| SYSb004659 | *torquatus* | TOR | Hefei, Anhui, China | 117.23 | 31.82 | Muscle | ♂ |  |  |  |  |  |  |  |  |  |
| SYSb004835 | *torquatus* | TOR | Hefei, Anhui, China | 118.43 | 31.34 | Muscle | ♂ |  |  |  |  |  |  |  |  |  |
| SYSb004656 | *torquatus* | TOR | Nanchang, Jiangxi, China | 115.97 | 28.40 | Muscle | ♀ |  |  |  |  |  |  |  |  |  |
| SYSb004657 | *torquatus* | TOR | Nanchang, Jiangxi, China | 115.97 | 28.40 | Muscle |  |  |  |  |  |  |  |  |  |  |
| SYSb004830 | *torquatus* | TOR | Shaoguan, Guangdong, China | 114.07 | 24.95 | Muscle | ♀ |  |  |  |  |  |  |  |  |  |
| SYSb004794 | *torquatus* | TOR | Yancheng, Jiangsu, China | 120.58 | 33.40 | Muscle | ♀ |  |  |  |  |  |  |  |  |  |
| SYSb004541 | *torquatus* | TOR | Yongshun, Hunan, China | 109.85 | 29.00 | Muscle |  |  |  |  |  |  |  |  |  |  |
| SYSb004602 | *vlangalii* | VLA | Dulan, Qinghai, China | 98.10 | 36.30 | Muscle | ♂ |  |  |  |  |  |  |  |  |  |
| SYSb004978 | *vlangalii* | VLA | Golmud, Qinghai, China | 94.93 | 36.41 | Muscle |  |  |  |  |  |  |  |  |  |  |
| SYSb004979 | *vlangalii* | VLA | Golmud, Qinghai, China | 94.93 | 36.41 | Muscle |  |  |  |  |  |  |  |  |  |  |
| SYSb004980 | *vlangalii* | VLA | Golmud, Qinghai, China | 94.93 | 36.41 | Muscle |  |  |  |  |  |  |  |  |  |  |
| SYSb004983 | *vlangalii* | VLA | Qaidam Basin, Qinghai, China | 101.50 | 36.65 | Muscle | ♂ |  |  |  |  |  |  |  |  |  |
| SYSb004984 | *vlangalii* | VLA | Qaidam Basin, Qinghai, China | 101.50 | 36.65 | Muscle | ♂ |  |  |  |  |  |  |  |  |  |
| SYSb004985 | *vlangalii* | VLA | Qaidam Basin, Qinghai, China | 101.50 | 36.65 | Muscle | ♂ |  |  |  |  |  |  |  |  |  |
| SYSb004986 | *vlangalii* | VLA | Qaidam Basin, Qinghai, China | 101.50 | 36.65 | Muscle | ♂ |  |  |  |  |  |  |  |  |  |
| SYSb005039 | *vlangalii* | VLA | Wulan, Qinghai, China | 98.48 | 36.93 | Muscle |  |  |  |  |  |  |  |  |  |  |
| SYSb006734 | *vlangalii* | VLA | Wulan, Qinghai, China | 98.48 | 36.93 | DNA |  |  |  |  |  |  |  |  |  |  |
| SYSb001824 | Outgroup (*Phasianus versicolor )* | VER | Japan | 143.52 | 43.81 | Feather | ♂ |  |  |  |  |  |  |  |  |  |
| SYSb001826 |  | VER | Japan | 143.52 | 43.81 | Feather | ♂ |  |  |  |  |  |  |  |  |  |
| SYSb001827 |  | VER | Japan | 143.52 | 43.81 | Feather | ♂ |  |  |  |  |  |  |  |  |  |
| SYSb001835 |  | VER | Japan | 143.52 | 43.81 | Feather | ♀ |  |  |  |  |  |  |  |  |  |
| SYSb002253 |  | VER | Japan | 143.52 | 43.81 | Feather |  |  |  |  |  |  |  |  |  |  |
| SYSb002256 |  | VER | Japan | 143.52 | 43.81 | Feather |  |  |  |  |  |  |  |  |  |  |
| SYSb002257 |  | VER | Japan | 143.52 | 43.81 | Feather |  |  |  |  |  |  |  |  |  |  |
| SYSb002258 |  | VER | Japan | 143.52 | 43.81 | Feather |  |  |  |  |  |  |  |  |  |  |
| SYSb002259 |  | VER | Japan | 143.52 | 43.81 | Feather |  |  |  |  |  |  |  |  |  |  |
| SYSb002262 |  | VER | Japan | 143.52 | 43.81 | Feather |  |  |  |  |  |  |  |  |  |  |

**Table S2** Primers sequences used for the amplification of two mitochondrial loci and seven nuclear introns.

| **Gene** | **Region** | **Length** | **Primer (5'→3')** | **reference** |
| --- | --- | --- | --- | --- |
| *CR* | control region of mitochondria | ~1000 | L16757: AGGACTACGGCTTGAAAAGC;  H1259: CATCTTGGCATCTTCAGTGCC | (Randi et al., 2000) |
| *cyt b* | Cytochrome of mitochondria | ~1000 | L14995: CTCCCAGCCCCATCCAACATCTCAGCATGATGAAACTTCG;  H16065: CTAAGAAGGGTGGAGTCTTCAGTTTTTGGTTTACAAGAC | (Kimball, Braun, Zwartjes, Crowe, & Ligon, 1999) |
| *AldB* | Aldolase b intron 6, nuclear Z-chromosome | ~600 | AldB6F: GAGCCAGAAGTCTTACCTGAYGG;  AldB7R: CAGCTGTCACCATGTTNGG | (Kimball et al., 2009) |
| *Bfibex* | Beta-fibrinogen Intron 4, nuclear autosomal | ~700 | BfibexF: CACGCCATATAGAGTATACTGTGACA;  BfibexR: AACACTACCATCCTGGCGATTCTGAA |  |
| *CLTC* | Clathrin heavy polypeptide intron 6, nuclear autosomal | ~1000 | CLTCF: CAGAATCCTGATCTAGCTTTACGAATGGC;  CLTCR: CATTTCTCCAGAAGTTGTTTGCGTCC |  |
| *DCoH* | Dimerization cofactor of HNF1 intron 3, nuclear autosomal | ~650 | DCOH3F: AGGCCTGGCTTCATGAC;  DCOH4R: GATAAACCYGTGCARTCYTGGGTGCT |  |
| *HMG17* | High mobility group 17 intron, nuclear autosomal 2 | ~650 | HMG172F: GCTGAAGGAGATACCAARGGCGA;  HMG174R: CTTTGGAGCTGCCTTTTTAGG |  |
| *SerpinC* | Serpin peptidase inhibitor clade C intron 5, nuclear autosomal | ~700 | SerpinCF: GTTCGCTTTGATAAACTTCCAGG;  SerpinCR: GGTGATTTGGTTGAGNATGTC |  |
| *OvoG* | Ovomucoid intron G, nuclear autosomal | ~500 | OVOGF: CAAGACATACGGCAACAARTG;  OVOGR: GGCTTAAAGTGAGAGTCCCRTT | (Armstrong et al., 2001) |

**Table S3** PCR amplification protocols for the analysed markers.

| **Reaction system** | | **Temperature system** | | | |
| --- | --- | --- | --- | --- | --- |
| **Reagents** | **volume** | **Stages** | **Temperature** | **Duration** | **Cycles** |
| PCR buffer (2×) | 5 μl | Initial denaturation | 94°C | 2 min |  |
| dNTP (2.5 mM) | 2 μl | Denaturation | 94°C | 30s | 10 |
| forward primers (10 μM) | 0.5 μl | Annealing | 58°C, decreasing 1°C per cycle | 30s |  |
| reverse primers (10 μM) | 0.5 μl | Extension | 72°C | 90s |  |
| *Taq* polymerase (5 unit/μl) | 0.2 μl | Denaturation | 94°C | 30s | 35 |
| template DNA (30-70 ng) | 1 μl | Annealing | 48°C | 30s |  |
| ddH_2_O | 0.8 μl | Extension | 72°C | 90s |  |
|  |  | Final extension | 72°C | 10min |  |

**Table S4** Polymorphic information of each locus. (*S,* number of segregating sites; *G+C*, GC content; *h*, number of haplotypes; *Hd*, Haplotype diversity; *π*, nucleotide diversity; S.D., Standard Deviation; *k*, Average number of nucleotide differences).

| **Locus** | **Length** | ***S*** | ***G+C*** | ***h*** | ***Hd*** | **S.D.** | ***π*** | **S.D.** | ***k*** |
| --- | --- | --- | --- | --- | --- | --- | --- | --- | --- |
| *CR* | 498 bp | 44 | 0.488 | 63 | 0.969 | 0.002 | 0.02135 | 0.00047 | 7.82 |
| *cyt b* | 690 bp | 47 | 0.484 | 41 | 0.914 | 0.007 | 0.00880 | 0.00045 | 5.88 |
| mtDNA | 1188 bp | 94 | 0.484 | 86 | 0.977 | 0.002 | 0.01323 | 0.00043 | 13.70 |
| *AldB* | 540 bp | 14 | 0.378 | 12 | 0.352 | 0.030 | 0.00129 | 0.00014 | 0.70 |
| *Bfibex* | 612 bp | 14 | 0.343 | 16 | 0.688 | 0.017 | 0.00283 | 0.00010 | 1.71 |
| *CLTC* | 905 bp | 28 | 0.471 | 46 | 0.903 | 0.006 | 0.00291 | 0.00008 | 2.62 |
| *DCoH* | 591 bp | 32 | 0.492 | 48 | 0.906 | 0.008 | 0.00552 | 0.00018 | 3.18 |
| *HMG17* | 697 bp | 46 | 0.479 | 52 | 0.913 | 0.007 | 0.00393 | 0.00015 | 2.73 |
| *OvoG* | 459 bp | 15 | 0.475 | 36 | 0.890 | 0.007 | 0.00657 | 0.00022 | 2.98 |
| *SerpinC* | 617 bp | 28 | 0.407 | 43 | 0.855 | 0.013 | 0.00400 | 0.00018 | 2.45 |

**Table S5** Estimates of genetic distances (*D*) and substitution rates of each locus.

| **Locus** | **Substitution Model** | ***D*** | **S.D.** | **distance ratio** | **substitution rate (per site per million year)** | **substitution rate (per locus per year)** | **interval of substitution rate （per locus per year）** | |
| --- | --- | --- | --- | --- | --- | --- | --- | --- |
| *CR* | GTR+I+G | 0.0228 | 0.0036 | 2.6512 | 0.0315 | 1.5711E-05 | 1.32306E-05 | 1.81921E-05 |
| *cyt b* | HKY+I+G | 0.0086 | 0.0017 | 1.0000 | 0.0119 | 8.2110E-06 | 6.5879E-06 | 9.8341E-06 |
| mtDNA | GTR+I+G | 0.0144 | 0.002 | 1.6744 | 0.0199 | 2.3672E-05 | 2.03839E-05 | 2.69593E-05 |
| *AldB* | HKY+I | 0.0011 | 0.0004 | 0.1279 | 0.0015 | 8.2193E-07 | 5.23047E-07 | 1.12081E-06 |
| *Bfibex* | HKY+I | 0.0022 | 0.0012 | 0.2558 | 0.0030 | 1.8630E-06 | 8.46837E-07 | 2.87925E-06 |
| *CLTC* | HKY+I | 0.0025 | 0.0009 | 0.2907 | 0.0035 | 3.1307E-06 | 2.00363E-06 | 4.25771E-06 |
| *DCoH* | K80+I+G | 0.0038 | 0.0011 | 0.4419 | 0.0053 | 3.1076E-06 | 2.208E-06 | 4.00712E-06 |
| *HMG17* | HKY+I | 0.003 | 0.001 | 0.3488 | 0.0042 | 2.8934E-06 | 1.92891E-06 | 3.85781E-06 |
| *OvoG* | SYM+I+G | 0.0037 | 0.0014 | 0.4302 | 0.0051 | 2.3500E-06 | 1.46079E-06 | 3.23915E-06 |
| *SerpinC* | HKY+I | 0.0032 | 0.0009 | 0.3721 | 0.0044 | 2.7320E-06 | 1.96364E-06 | 3.5004E-06 |

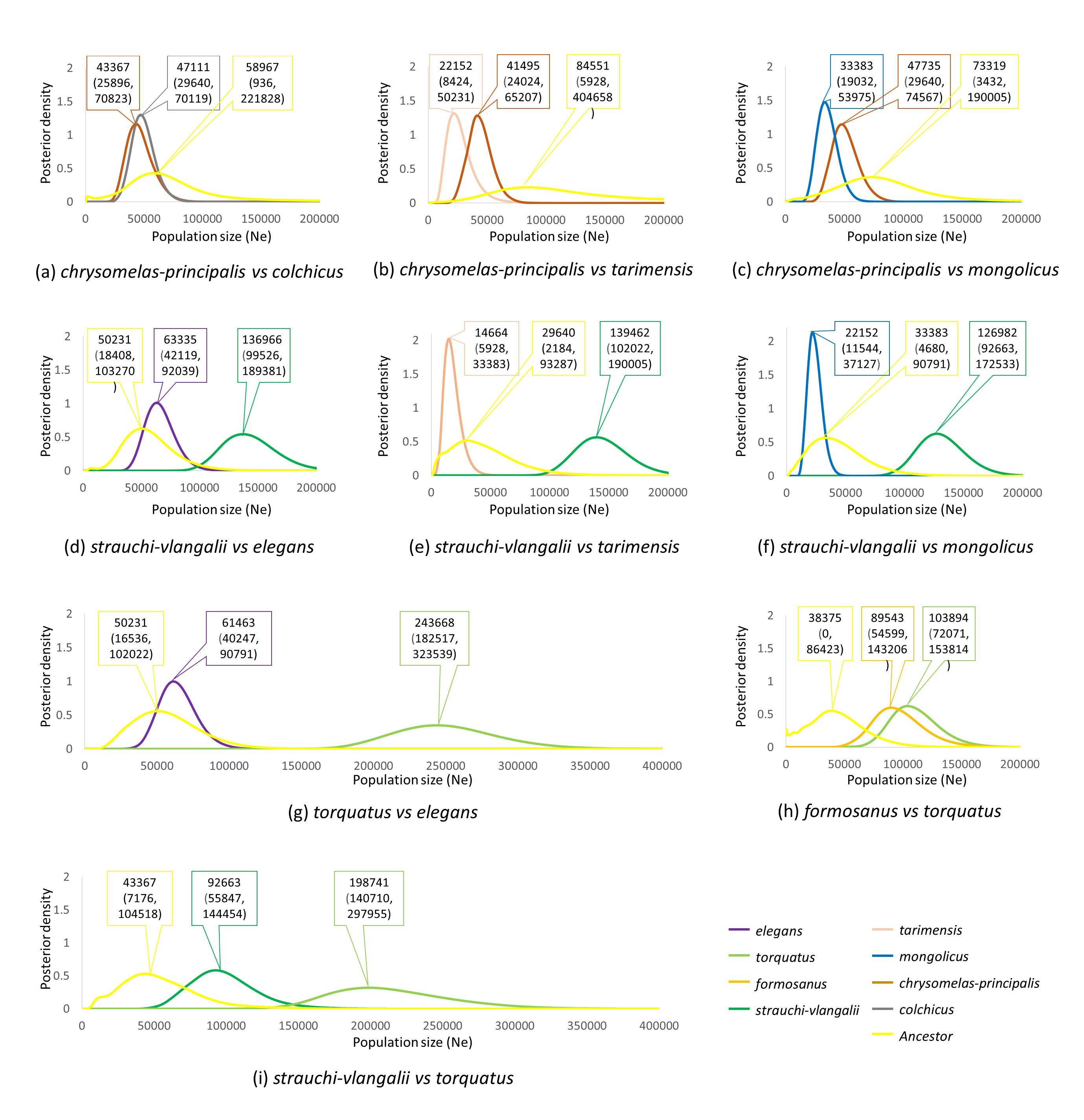

**Figure S1** Posterior probability distributions of effective population sizes between parapatric populations of the common pheasant based on concatenated data estimated in Ima2. Values indicate the highest values of the posterior distribution and the 95% highest posterior density interval.
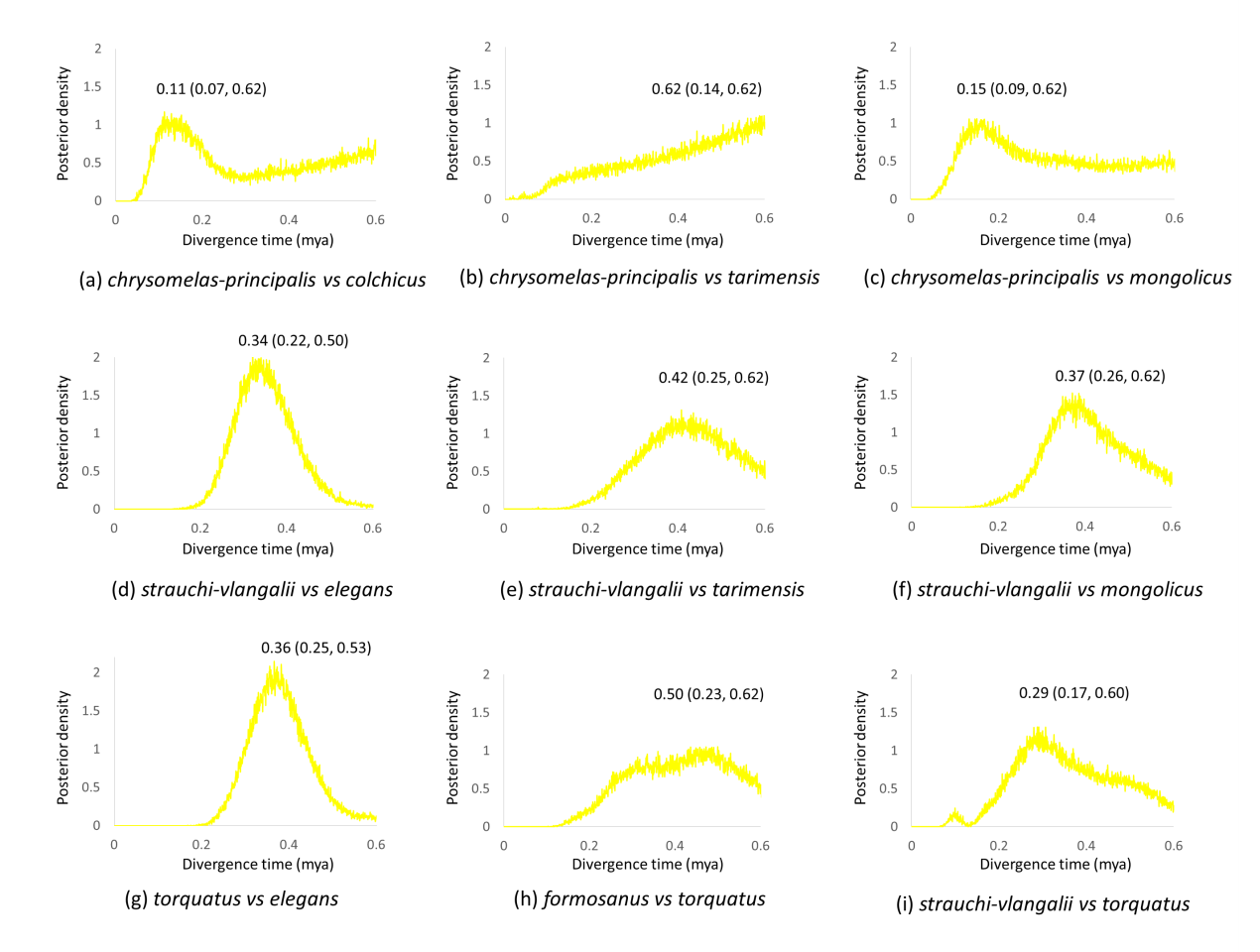

**Figure S2** Posterior probability distributions of divergence time estimate between parapatric populations of the common pheasant based on concatenated data estimated in Ima2. Values indicate the highest values of the posterior distribution and the 95% highest posterior density interval.
